# Visceral signaling of post-ingestive malaise directs memory updating in Drosophila

**DOI:** 10.1101/2025.10.21.683769

**Authors:** Bhagyashree Senapati, Christoph D. Treiber, Scott Waddell

## Abstract

Consolidation is a time when labile memories transition to a stable form. Malaise learning in *Drosophila* reveals consolidation to also permit memory updating. Flies taught to associate one of two odors with toxin-tainted sugar initially express conditioned odor approach, that following consolidation switches to avoidance. Behavioral reversal emerges from dopaminergic update of parallel memories for the two trained odors. Differential serotoninergic modulation of specific aversive and rewarding dopaminergic neuron subtypes permits post-ingestive intoxication to suppress consolidation of initial odor-sugar memory and simultaneously invert reward memory plasticity into “safety” memory for the odor experienced without food. Fat body release of the Toll-ligand activating protease modSP, and resilience factor Turandot A, instruct malaise updates by triggering autocrine Toll signaling in the same brain dopaminergic neurons that form and consolidate initial sugar memory. This neural mechanism overcomes the credit assignment problem of delayed post-ingestive reinforcement by updating earlier memories of the trained odors.

## Introduction

Much of our experience has conscious and subconscious elements. This is particularly apparent with feeding events, where hedonic orosensory properties, such as the taste, smell, appearance and texture of food are followed by initially subconscious post-ingestive effects, which can be positive (nutrition / satiety), negative (intoxication / malaise), or a combination of the two^1–5^. Remembering the full content of these experiences therefore requires neural mechanisms that bridge time between the temporally separate properties, to appropriately link them as parts of the same memory.

Studies of classical conditioning defined temporal rules for learning associations. Sensory stimuli that either briefly precede reinforcement, or are temporally contingent with it, are most efficiently learned ^6^. Moreover, the efficiency of associative learning sharply declines with increasing time between stimulus presentation and reinforcer delivery – the inter stimulus interval (ISI). Optimal ISIs vary between tasks and animals but they typically range from milliseconds to seconds. For example, eye-blink conditioning in humans operates between 100 ms and 1 s ^7^, in rabbits 200-500 ms is optimal ^8^, whereas aversive olfactory conditioning with shock reinforcement in *Drosophila* can tolerate an ISI of 15 s ^9–11^. Importantly, even this relatively long effective ‘trace’ period for classical conditioning is much too short to account for learning using post-ingestive reinforcement. Classic studies of Conditioned Taste Aversion (CTA) learning using poisoned bait ^12^, saccharin followed by γ-irradiation ^1^, or intragastric injection of malaise-inducing drug ^13^, established that the ISI between rats tasting food and experiencing sickness can be 1-3 hours. Importantly, a single CTA trial forms long term memory (LTM) that can last for months and even a year for rats surviving consumption of red squill. An early study in rats showed that CTA can also potentiate learning of an associated odor (taste potentiated odor aversion, TPOA), with odor producing a stronger conditioned aversion than the taste/flavor component ^14, 15^. CTA has been demonstrated in many mammals and birds and several invertebrates using ingestion of pathogenic bacteria or toxins, and sometimes using TPOA paradigms ^16–20^. Studying TPOA permits analyses of malaise memory that can be measured independently of the consummatory processes of tasting and ingesting food.

Cellular memory consolidation is operationally defined as a window of time during the first hours after training in which disruption of neuronal activity (or new transcription or protein synthesis) disrupts formation of stable long-term memory (LTM) ^21–25^. In *Drosophila* cellular consolidation to LTM occurs in close temporal relationship with a process that resembles ‘systems consolidation’, whereby the neuronal substrate required for memory expression transitions between parallel populations of mushroom body (MB) Kenyon Cells; with the γ KCs directing short-term appetitive memory (STM) and the αβ KCs being essential for LTM ^26–29^. The timing of these LTM-relevant processes suggests that the consolidation window overlaps with the arrival of post-ingestive reinforcement. Consistent with this notion, LTM formation in honeybees and *Drosophila* requires sugar ingestion ^30^, and for the sugars to be nutritious ^31–33^ with one published exception ^34^.

Studies in *Drosophila* have established that subdivision of the dopaminergic system supports a variety of forms of aversive and reward learning ^32, 35–50^. Distinct groups of DANs primarily form valence-specific memories by directing depression of synaptic connections between odor activated KCs and specific mushroom body output neurons (MBONs) ^51–57^. In the context of feeding-reinforced learning, different DANs appear to provide parallel representations of pre-ingestive taste (those to the γ4 compartment of γ KCs and β’2 of α’β’ KCs) and post-ingestive nutritional properties of sugars (those to the α1 compartment of αβ KCs) ^40, 41, 58^. Moreover, some DANs show increased activity after training that depends on the nutritious value of the sugar (PPL1-γ1pedc ^33^), and their output and presumably released dopamine, is required at this time for consolidation of appetitive LTM ^33, 58^.

Flies can also learn with water reward and these teaching signals and thirst-dependent motivational control appears to involve different DANs to those involved in sugar and hunger-dependent processes ^43, 44, 59^. In addition, a more recent study in the mouse demonstrated that different DANs track specific nutrients and fluids at different stages of ingestion ^60^. Notably, specific DANs selectively respond to systemic hydration within 10 minutes of fluid intake, allowing mice to learn flavor preferences based on a fluid’s rehydrating effect. Thus, in both flies and mice distinct subsystems of the larger dopaminergic system encode separable aspects of ingestion and internal nutritional status.

Here we investigated mechanisms of post-ingestive malaise/intoxication learning in *Drosophila* using a TPOA learning paradigm ^14, 15^ ^61^. Flies trained by pairing one of two odors with sucrose laced with the tasteless toxin amygdalin initially express conditioned approach that over time switches to aversion. Delayed reinforcement of post-ingestive malaise blocked consolidation of the sugar associated odor memory and formed a ‘safety’ memory for the other odor. These memory updates are permitted during the consolidation window by differential serotonergic modulation of five distinct subpopulations of DANs, which can be seen to direct both initial sugar-evoked plasticity and malaise-evoked reversals of these engrams. These specific DANs respond to malaise through autocrine Toll signaling, that depends upon humoral release from the fat body of an initiating autocatalytic serine protease and the Turandot A resilience factor.

## Results

### Flies learn to associate odors with malaise

To investigate neural mechanisms of post-ingestive malaise memory formation we replicated a learning protocol established in the honeybee, where odor is paired with feeding of sucrose laced with amygdalin, a tasteless cyanogenic glycoside ^61^, which induces illness in honeybees and humans ^62, 63^.

We first tested whether Drosophila are able to taste amygdalin. Starved flies showed no preference between arms of a T-maze containing either sucrose or sucrose plus amygdalin (SA) (Figure 1A). In contrast, flies robustly avoided sucrose mixed with bitter-tasting quinine (SQ) versus sucrose. A similar inability to taste amygdalin was evident when it was applied to the fly’s proboscis. Whereas inclusion of quinine inhibited sucrose-evoked proboscis extension, 0.6% amygdalin had no effect (Figure 1B). Lastly, amygdalin did not inhibit consumption of sucrose (Figure 1C), or abdominal distension (Figure 1D), a proxy for food ingestion. We therefore concluded that flies cannot taste amygdalin when presented at 0.6% with sucrose. We also tested whether flies displayed signs of malaise by tracking the spontaneous movement of individuals following toxin ingestion. Flies were largely immobile 15 min after toxin feeding (they however express memory performance, see below) but had significantly recovered 24 h later (Figure S1).

**Figure 1.**
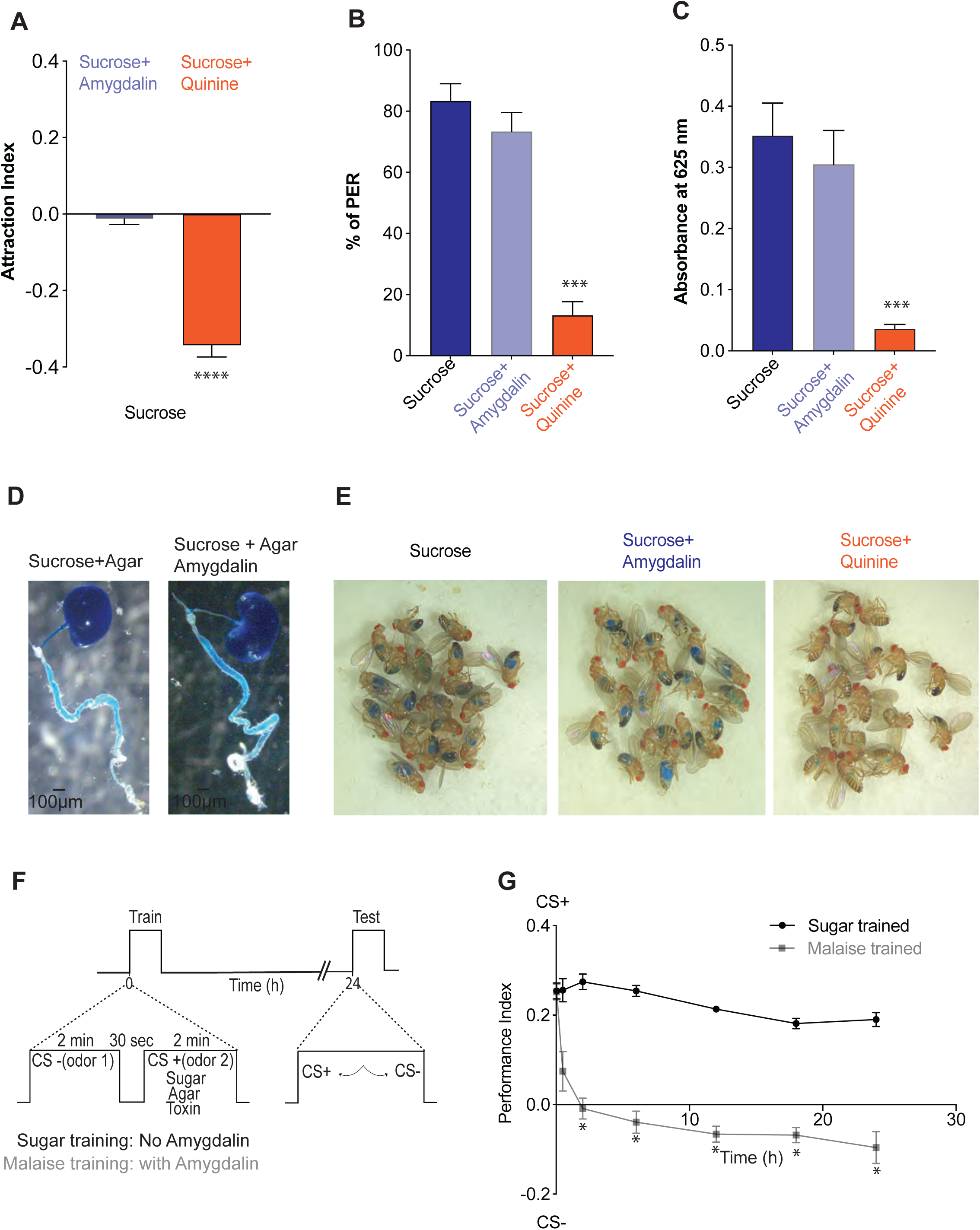
Flies learn to associate odors with post-ingestive malaise. (A) Naïve preference of starved flies between sucrose and sucrose +0.6% amygdalin (SA, blue), between sucrose and sucrose +0.6% quinine (SQ, orange). (B) Likelihood of eliciting Proboscis Extension Reflex (PER) in starved flies presented with sucrose, SA or SQ. (C) Food consumption is robustly inhibited by including quinine, but not amygdalin, in sucrose. (D) Distension of the gut and crop in flies fed sucrose or SA containing blue food dye. (E) Ingestion of blue food dye is apparent as colored abdomens in flies fed with sucrose or SA, but not SQ. (F) Schematic of regular sucrose rewarded, and malaise reinforced olfactory learning protocols. (G) Memory performance timelines of sucrose trained (black) or malaise trained (grey) flies at 2, 30 min, and 2, 6, 12, 18 and 24 h. Performance of the two groups was different at every time point except 2 min after training. Figure A data was compared with one sample t-test and Wilcoxon test. Figure B, C, & G were compared using one-way ANOVA with Tukey’s test; n ≥ 10. Data are mean ± Standard Error of Mean (SEM). Asterisks denote significant difference ^∗^*p* < 0.05, ^∗∗^*p* < 0.01, ^∗∗∗^*p* < 0.001, ^∗∗∗∗^*p* < 0.0001.

We next trained food-deprived flies using the standard sucrose-rewarded olfactory memory protocol ^27, 64^, or with an adapted ‘malaise’ variant where the second of two odor presentations was paired with sucrose containing 0.6% amygdalin (SA) (See Figure S2A for protocol optimization). Memory performance was subsequently measured by testing the flies’ preference between the previously food-paired odor (conditioned stimulus plus, CS+) and the previously unpaired odor (conditioned stimulus minus, CS-).

As expected, flies trained with odor and sucrose showed a persistent approach to the previously rewarded odor (Figure 1G). In contrast, SA trained flies initially showed strong approach to the previously food-paired odor which rapidly declined within the first 120 min after training and converted to avoidance by 6 h that persisted for at least 24 h. This SA memory timeline is consistent with flies forming a sugar reward memory that is cancelled by delayed post-ingestive aversive reinforcement. Importantly, flies did not show 24 h odor preference if SA was paired with the first of the two odor presentations during training, or if flies were differentially conditioned by pairing one odor with sucrose and the other with SA (Figure S2B). In addition, flies trained with the pre-ingestive taste-dependent SQ protocol did not show conditioned approach at any time (Figure S2B), consistent with published reports of training with sugar laced with a strongly bitter insect repellent DEET ^54^ (Figure S2C). We confirmed that the observed dynamic of SA memory did not result from impaired odor acuity, reduced motivation (locomotion or feeding), or general lethargy from sickness (Figure S2D, S2E & S2F).

### Opposing valence dopaminergic neurons are required for malaise learning

Given the established roles for discrete types of dopaminergic neurons in reward and aversive learning in insects and mammals, we tested their involvement in malaise learning ^32, 35–50^. The dominant negative temperature sensitive UAS-*Shibire*^ts^^1^ transgene ^65^ was expressed in either rewarding (R58E02-GAL4) or aversively reinforcing (TH-GAL4) DANs (Figure 2A). We then blocked DAN output during learning by training the flies at restrictive 33°C. Flies were returned to permissive 25 °C immediately after training and subsequently tested for 24 h malaise memory (Figure 2B). Blocking either rewarding or aversive DANs abolished formation of malaise memory, suggesting that malaise learning engages DANs of opposing valence.

**Figure 2.**
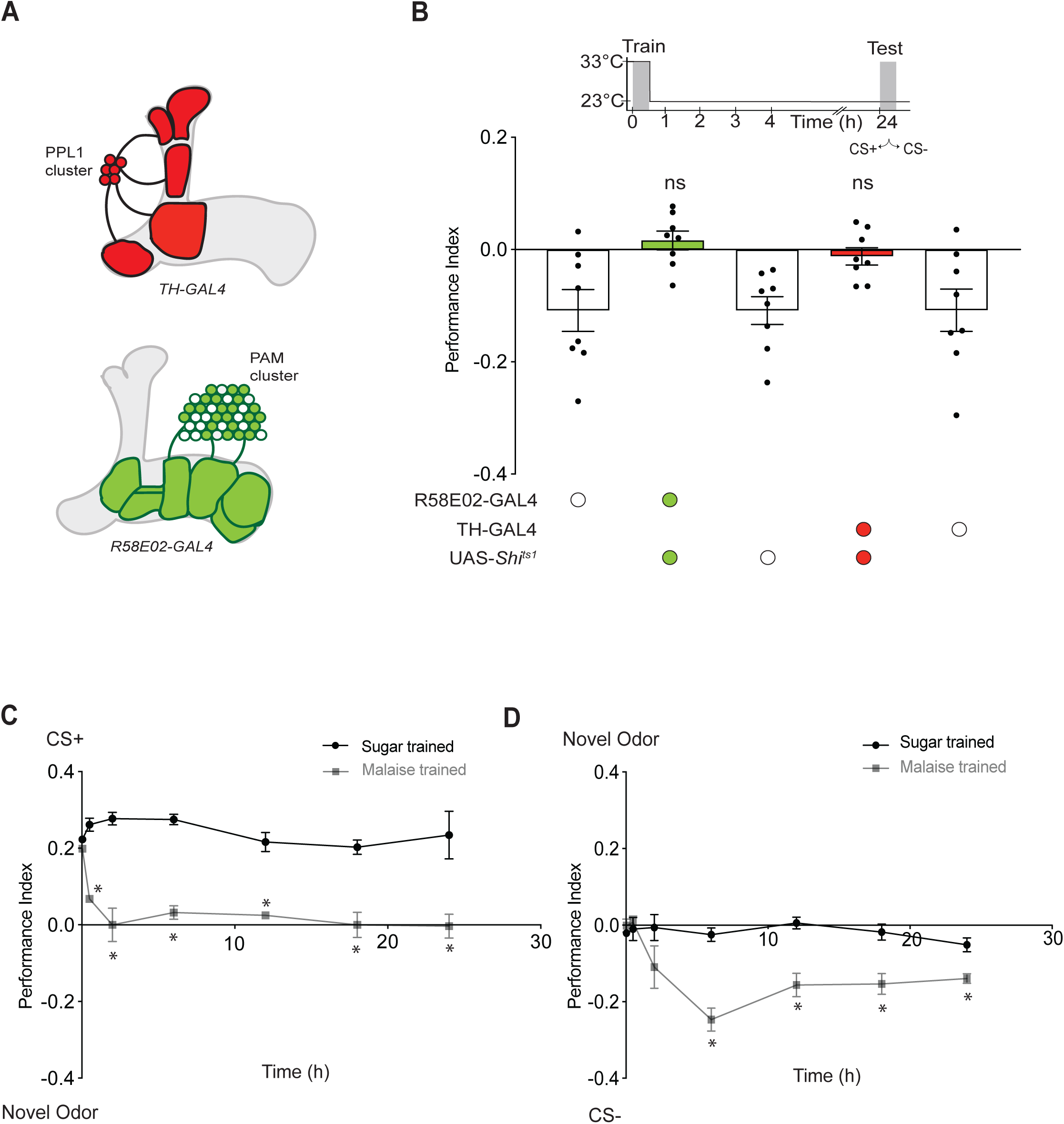
Opposing valence dopaminergic neurons are required for malaise learning. (A) Schematic of PPL1 (TH-GAL4) and PAM (R58E02-GAL4) dopaminergic neurons innervating the mushroom body (MB). The MB is bilaterally symmetrical but only the fly’s right-hand hemisphere is shown. (B) Upper panel: temperature shifting protocol imposed on training and testing timeline. Lower panel: blocking output from either the PAM or PPL1 DANs during training using UAS-*Shi^ts^* abolishes 24 h malaise memory. (C) Timelines of CS+ memory measured v novel odor for both sugar trained, and malaise trained flies at 2 and 30 min, 2, 6, 12, 18 and 24 h. Performance of the two groups was different at every time point except 2 min after training. (D) Timelines of CS-memory measured v novel odor for sugar trained and malaise trained flies. Memory performance of the two groups is statistically different from 6 h onwards. Data in Figure B was compared with one sample t and Wilcoxon tests. Data in Figure C & D were compared using one-way ANOVA with Tukey’s test, n ≥ 8; Data are mean ± SEM; dots represent individual data points, which correspond to independent groups of approximately 200 flies. Asterisks denote significant differences: ^∗^*p* < 0.05, ^∗∗^*p* < 0.01, ^∗∗∗^*p* < 0.001, ^∗∗∗∗^*p* < 0.0001.

The observed conversion from approach to avoidance could result from a developing avoidance memory for the CS+ and/or learning that the CS-was not feeding/malaise associated. Therefore, rather than testing flies for preference between CS+ and CS-odors, they were tested for preference between CS+ versus a novel odor or CS-versus novel odor. We again compared the malaise memory timelines to those of flies trained using sugar reward. (Figure 2C & D). Sugar-trained flies maintained a preference over 24 h for the previously rewarded CS+ over novel odor (Figure 2C). In contrast, malaise-trained flies initially showed CS+ approach but by 2 h after training no odor preference was apparent, suggesting the CS+ approach memory was quickly lost. Moreover, malaise trained flies showed a delayed emergence of a CS-approach memory (Figure 2D), which was not formed in sugar-trained flies. We noted that CS-approach measured at 2 and 6 h versus novel odor was greater than that observed at the same time versus CS+ (compare Figure 1G and Figure 2D). We therefore conclude that malaise value forms parallel CS+ and CS-memories and that the flies might also learn to be generally wary of odors they have not previously experienced.

### Delayed DPM neuron action is required for malaise update memories

Output from the serotonergic Dorsal Paired Medial (DPM) neurons is required during the first hour after training to consolidate sugar reward memories ^27, 66–68^. We therefore tested whether similarly timed DPM neuron activity was required for malaise memory formation. We first used expression of UAS-*Shi*^ts^^1^ to temporarily block DPM neuron output and assessed the effects on CS-approach memory. Blocking DPM for 1 h during and after training significantly impaired malaise-induced CS-approach memory measured 6 h after training, whereas later block between 1-3 h, or 2-4 h, after training had no effect (Figure 3A). To refine the 1 h time point, we inhibited DPM neurons using the green light-gated anion channel GtACR1 ^69^, which permits greater temporal resolution than *Shi*^ts^^1^ (Figure 3B). These experiments identified DPM neurons to be required 15-30 min after training to form malaise-induced CS-approach memory, that we then replicated using *Shi*^ts^^1^ (Figure 3C). Strikingly, inhibiting DPM neurons 15-30 min after training also reinstated the malaise-driven ‘loss’ of CS+ approach memory (Figure 3D). The DPM blocking experiments therefore demonstrate that 15-30 min after training is a critical time for malaise information to both suppress sugar-rewarded CS+ approach memory and form a CS-approach memory. In addition, since the 15-30 min consolidation block restores 24 h CS+ approach (Figure 3D), this DPM manipulation must spare the consolidation to LTM driven by the post-ingestive nutrient value of sucrose.

**Figure 3.**
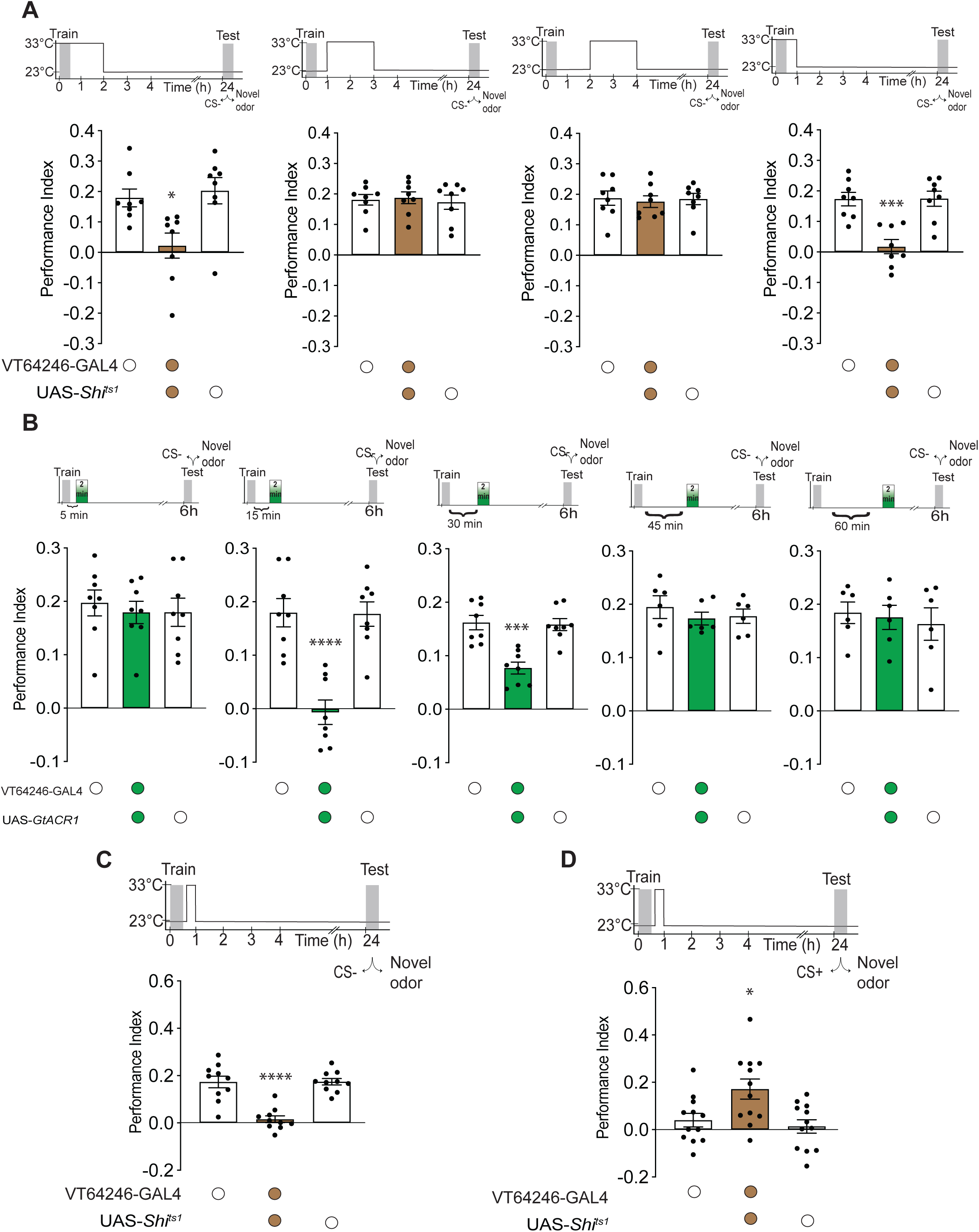
Delayed DPM neuron action is required for malaise update memories. (A) Upper panels, temperature shifting protocols imposed on training and testing timelines. Lower panels, 24 h CS-memory performance measured versus novel odor, following UAS-*Shi^ts^* mediated block of DPM neurons (VT46246-GAL4) for 0-2, 1-3, 2-4 or 0-1 h after training. A significant difference within the controls was only observed with block 0-1 or 0-2 h. (B) Upper panels, illumination protocols for transient inactivation of DPM neurons at various times after training and before testing imposed on training and testing 6 h CS-memory performance measured versus novel odor. Lower panels, 2 min optogenetic silencing of DPM neurons, using green light to trigger GtACR1, impairs memory if delivered 15 or 30 min after training, but not at 5, 45 or 60 min. (C and D) Upper panels, temperature shifting protocols imposed on training and testing timelines. (C) Lower panel, blocking output for 15-45 min following training from DPM neurons using UAS-*Shi^ts^* impairs CS-memory measured vs novel odor. (D) Lower panel, blocking output for 15-45 min following training from DPM neurons using UAS-*Shi^ts^* restores CS+ approach memory measured vs novel odor. Data in Figures A-D were compared using one-way ANOVA with Tukey’s test, *n*=8; Individual data points represent independent groups of approximately 200 flies. Asterisks denote significant differences: ^∗^*p* < 0.05, ^∗∗^*p* < 0.01, ^∗∗∗^*p* < 0.001, ^∗∗∗∗^*p* < 0.0001.

### Different 5-HT receptors are required in specific DAN subtypes for malaise memories

The presynaptic terminals of DPM neurons lie exclusively within the neuropil of the MB lobes and base of the peduncles ^70, 71^. There they interact with the presynaptic terminals of DANs, dendrites of Mushroom Body Output Neurons (MBONs), the inhibitory APL neurons and the presynaptic terminals of KCs ^71^. Knowing aversive and appetitive DANs were required for malaise learning, we investigated whether serotonin (5-hydroxytryptamine, 5-HT) signaling might reveal the specific DAN types directing formation of CS+ and CS-update memories.

*Drosophila* have five 5-HT receptor subtypes; 5-HT1A, 5-HT1B, 5-HT2A, 5-HT2B, and 5-HT7 ^72–76^ and single-cell sequencing suggests that they are differentially expressed in DANs (Figure S4 and Treiber et al., in prep). We therefore first used transgenic UAS-RNA interference (RNAi) constructs to knockdown each 5-HT receptor in most rewarding DANs. These analyses revealed defective malaise-induced CS-memory in flies with 5-HT1A, 5-HT2A, and 5-HT7 loss-of-function (Figure S5). We subsequently performed DAN subtype specific RNAi experiments for these 3 receptors, which identified 5-HT7 to be required in PAM-β’2mp and PAM-γ3 DANs, and 5-HT1A in PAM-γ5 neurons, for malaise-induced CS-approach memory (Figure 4A & S6). We did not observe clear defects with DAN type-specific loss of 5-HT2A (data not shown).

**Figure 4.**
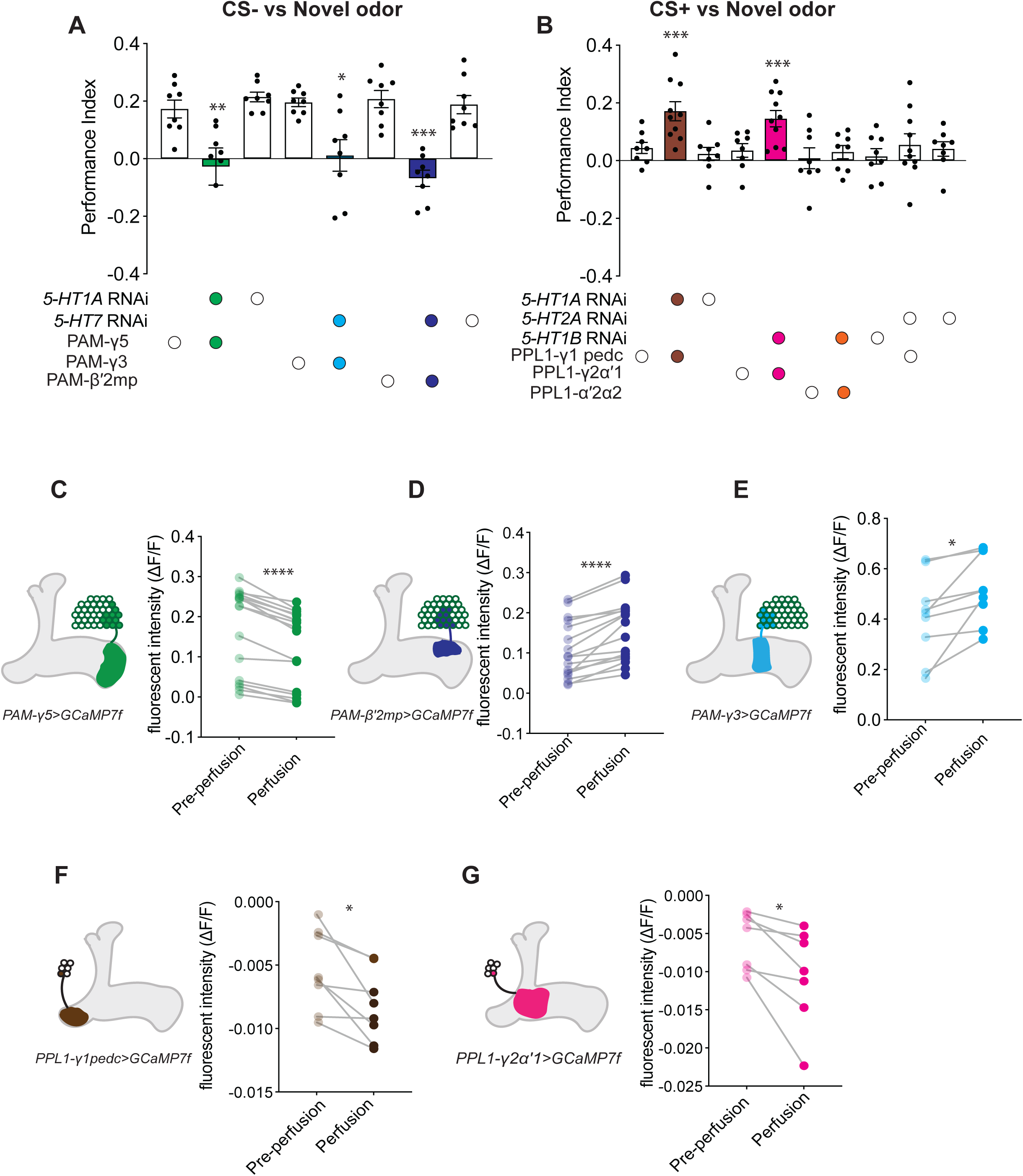
Differential 5-HT receptor directed modulation of DAN subtypes underlies malaise learning. (A) RNAi-mediated knockdown of 5-HT1A in γ5 PAM-DANs and 5-HT7 in γ3 or β’2mp PAM-DANs impairs malaise-induced CS-approach memory measured at 24 h v novel odor. (B) RNAi-mediated knockdown of 5-HT1A in γ1pedc and 5-HT1B in γ2α′1 PPL1 DANs restores CS+ approach memory measured at 24 h v novel odor. (C-G) Left, schematics of γ5, β’2mp γ3, γ2α’1 & γ1pedc DANs. Right, Calcium imaging values before and during perfusion of 5-HT (in presence of TTX) onto γ5, β’2mp, γ3, γ2 α’1 & γ1pedc DANs expressing UAS-*GCaMP7f*. Data in Figure (A and B) were compared using one sample t and Wilcoxon test. Data in figs (C-G) were compared using t tests (non-parametric tests) with Wilcoxon matched pairs signed rank test, n≥10. Figure (A, B), individual data points represent independent groups of approximately 200 flies. Data in Figure. (C-G), individual data points show each fly’s response. Asterisks denote significant differences: ^∗^*p* < 0.05, ^∗∗^*p* < 0.01, ^∗∗∗^*p* < 0.001, ^∗∗∗∗^*p* < 0.0001.

We also tested the roles of 5-HT receptors in specific types of aversive PPL1 DANs in the malaise-driven loss of CS+ approach memory. Strikingly, 24 h CS+ approach memory was retained in flies with RNAi knockdown of 5-HT1B in PPL1-γ2α’1 or PPL1-α’2α2 neurons, and of 5-HT1A in PPL1-γ1pedc neurons (Figure 4B). Involvement of PPL1 DANs is consistent with loss of CS+ approach performance resulting from formation of a competing parallel aversive malaise memory ^45, 77^ or in blocking the nutrient dependent consolidation of sugar rewarded LTM for the CS+ odor ^33^.

5-HT1A, 5-HT1B and 5-HT7 receptors are known to be differentially coupled to inhibitory Gi and stimulatory Gs proteins, respectively ^78^. To test how 5-HT receptors are coupled in the malaise learning-relevant DANs, we monitored their activity using DAN specific expression of the GCaMP7f calcium indicator. 100 μM 5-HT (in the presence of tetrodotoxin to block indirect neuronal input) was bath applied to exposed brains of flies expressing GCaMP7f in PAM-β’2mp, PAM-γ3, PAM-γ5 PPL1-γ1pedc or PPL1-γ2α’1 DANs (Figure 4C-G). 5-HT significantly decreased GCaMP7f fluorescence in PPL1-γ1pedc, PPL1-γ2α’1 and PAM-γ5 DANs (Figure 4F, 4G and 4C) but increased the signal in PAM-γ3 and PAM-β’2mp DANs (Figure 4D & 4E). These data suggest that the same DPM-released 5-HT, through differential 5-HTR coupling, inhibits some DAN subtypes while facilitating others to permit malaise-directed memory updates. The observed 5-HT evoked effects generally reflect the signaling modes expected for the 5-HT receptors identified in each DAN subtype by receptor loss-of-function, although in some cases they express multiple 5-HT receptors (Figure S4).

### Malaise memory requires differential DAN subtype activity during consolidation

To directly test the inferred modes of 5-HT-receptor action we temporally manipulated activity of the relevant DANs 15-45 min after malaise training, since that is the precise time in which both CS+ and CS-memories were observed to change (Figure S3). We expressed a UAS-transgene for either the heat-sensitive Ca^2+^ channel dTrpA1 ^79^ or Shi^ts^^1^ using DAN-specific control. These effector transgenes permit respective activation or block of the DANs using the same restrictive 33 °C temperature. Temporarily blocking output from PAM-β’2mp and PAM-γ3 DANs significantly impaired malaise-induced CS-memory, whereas no effect was observed with PAM-γ5 DAN block (Figure 5A). Conversely, dTrpA1-mediated activation of PAM-γ5 DANs 15-45 min post-training impaired malaise-induced CS-memory (Figure 5B), while no effect was observed with PAM-β’2mp or PAM-γ3 activation. These data are therefore consistent with malaise induced CS-memory formation requiring 5-HT receptor coupling that facilitates PAM-β’2mp and PAM-γ3 DANs while simultaneously permitting inhibition of PAM-γ5 DANs.

**Figure 5.**
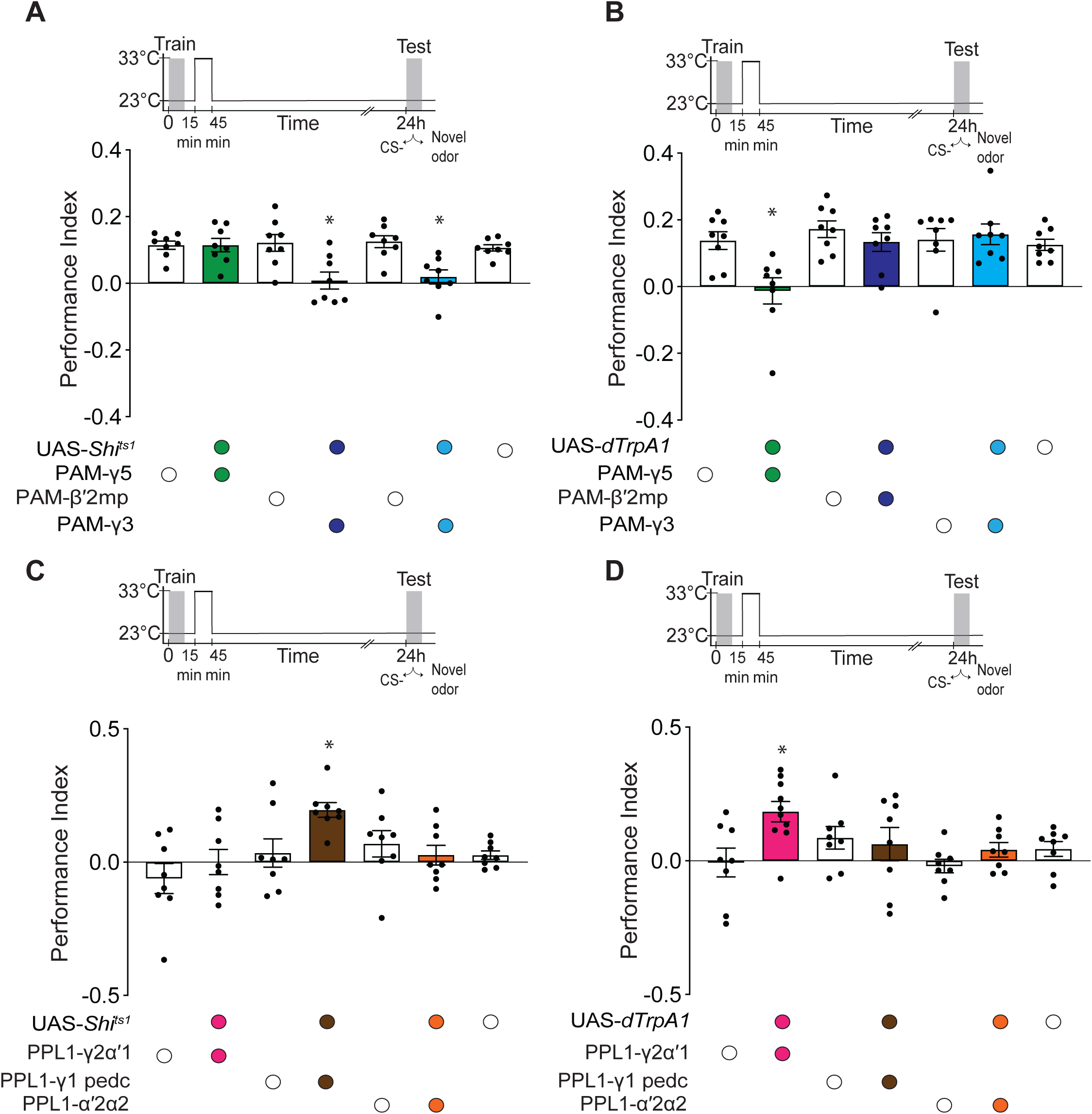
Malaise memory requires bidirectional DAN subtype activity during consolidation. (A and B) Upper panels, temperature shifting protocols imposed on training and testing timelines. (A) Lower panel, UAS-*Shi^ts^* mediated block of γ3 and β’2mp DANs for 30 min 15 min after training impairs malaise-induced CS-approach memory measured at 24 h v novel odor. (B) Lower panel, UAS-*dTrpA1* mediated activation of γ5 DANs for 30 min15 min after training impairs CS-approach memory measured at 24 h v novel odor. (C and D) Upper panels, temperature shifting protocols imposed on training and testing timelines. (C) UAS- *Shi^ts^* mediated block of PPL1- γ1pedc DANs for 30 min 15 min after training restores 24 h CS+ approach memory measured at 24 h v novel odor. (D) Lower panel, UAS-*dTrpA1* mediated activation of PPL1-γ2α′1 DANs for 30 min 15 min after training re-establishes CS+ approach memory measured at 24 h v novel odor. Data in Figures (B, C) were compared using one-way ANOVA with Tukey’s test, *n*=8; Data in Figures (F, G) were compared using one sample t and Wilcoxon test, n≥12. Individual data points represent independent groups of approximately 200 flies. Asterisks denote significant differences: ^∗^*p* < 0.05, ^∗∗^*p* < 0.01, ^∗∗∗^*p* < 0.001, ^∗∗∗∗^*p* < 0.0001.

To test for a similar mechanism underlying the malaise-directed update of CS+ memory, we independently controlled PPL1-γ2α’1, PPL1-α’2α2 and PPL1-γ1pedc DANs. Shi^ts^^1^-mediated block of PPL1-γ1-pedc DANs 15-45 min after training abolished the malaise-driven update of CS+ memory leading to re-emergence of 24 h CS+ approach (Figure 5C). Similarly timed block of PPL1-γ2α’1 or PPL1-α’2α’2 DANs had no effect on 24 h CS+ performance, which was indifferent from controls (zero). In contrast, dTrpA1-mediated activation of PPL1-γ2α’1 DANs produced re-emergence of 24 h CS+ approach, while PPL1-α’2α2 and PPL1-γ1pedc DAN activation had no significant effect (Figure 5D). Therefore, while these data corroborate roles for PPL1-γ1pedc and PPL1-γ2α’1 DANs in the abolition of CS+ approach memory, they suggest that 5-HT1A and 5-HT1B receptor directed modulation is more nuanced than just inhibiting function of the respective DANs during the consolidation window.

### Malaise reverses sugar reinforced KC-MBON plasticity

MB innervation of the relevant DANs dictates the identity of the KC-MBON connections that are modified by malaise learning. Importantly, retrieval-restricted blocking of β′2mp, γ5 and γ3/γ3β′1 MBONs impaired expression of malaise-updated CS- approach memories. Blocking the γ1pedc>αβ and γ2α’1 MBONs was inconsequential (Figure S9), presumably because they are required for expression of aversive memories. To visualize potential network manifestations of memory updating, we used in vivo calcium imaging of odor-evoked MBON responses 15 min, 2 h and 24 h after sugar-toxin training.

The β′2mp, γ5 and γ3/γ3β′1 MBONs exhibited a particularly striking time-dependent reversal of odor-specific plasticity. Both of the avoidance-directing β′2mp and γ5 MBONs showed relative depression of CS+ versus CS- responses 15 min after training, consistent with sugar-rewarded CS+ approach memory. However, their responses reversed to relative CS- depression at 24 h, representing CS- approach memory. (Figure 6A–D). The approach-directing γ3/γ3β′1 MBONs exhibited the opposite inversion from relative CS− versus CS+ depression at 15 min, to CS+ depression at 24 h (Figure 6E & 6F), which is consistent with γ3 DANs being suppressed by sucrose ^117^, and conversion of CS+ approach into CS- approach memory.

**Figure 6.**
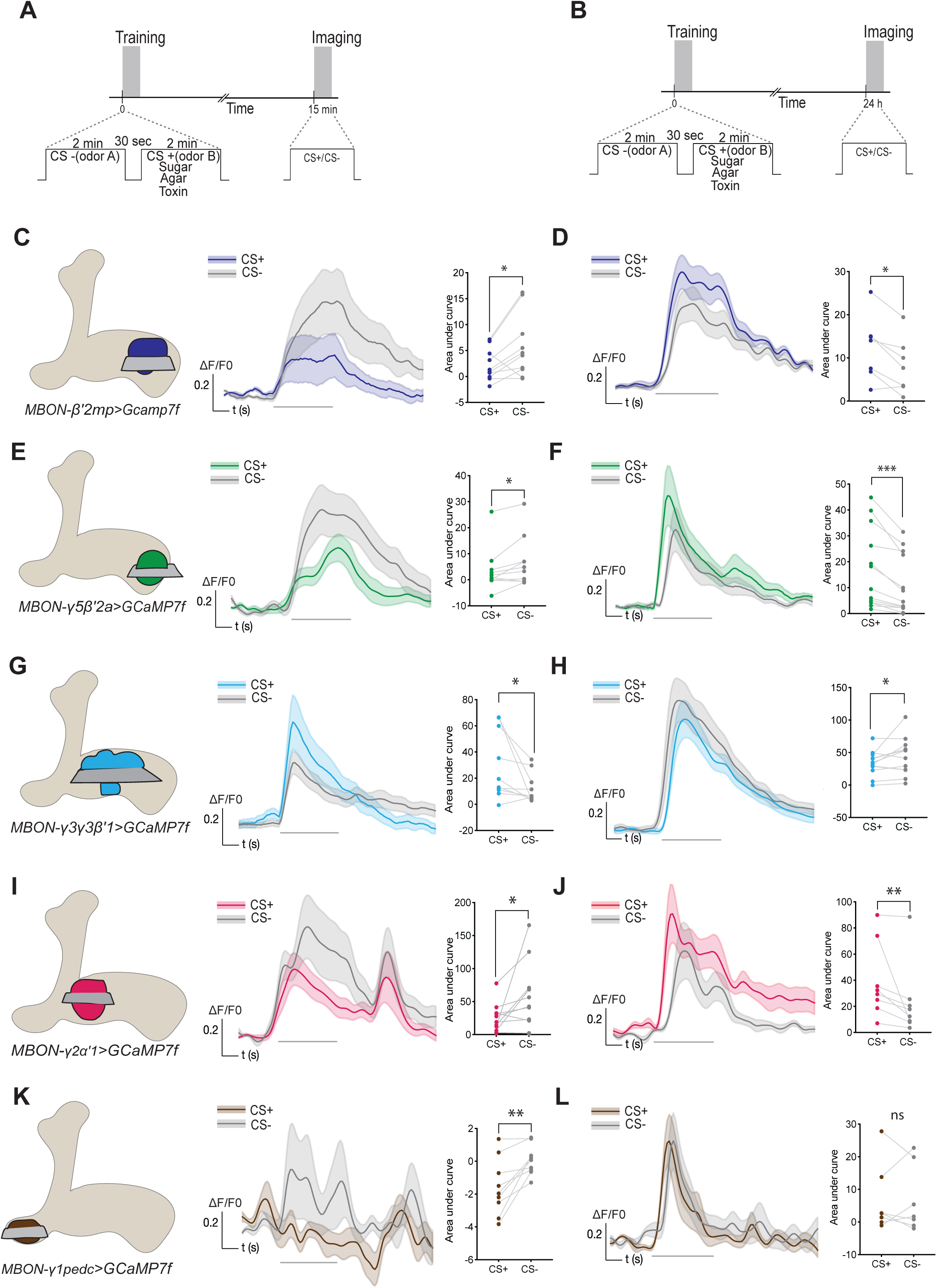
Malaise reverses sugar-reinforced KC–MBON plasticity. (A) Protocol for measuring odor-evoked responses in MBONs by in vivo imaging 15 min after training. (B) Protocol for measuring odor-evoked responses at 24 h. (C, E, G, I, K) Left: anatomy and imaging planes of MBONs β′2mp, γ5β′2a, γ3 with γ3β′1, γ2α′1, and γ1pedc>αβ. Middle: odor-evoked responses following malaise training at 15 min. Right: quantification of responses. (D, F, H, J, L) Left: Odor-evoked responses of the same MBONs at 24 h after training. Right: quantification of responses. (C) CS+ responses were significantly reduced versus those to CS- in β′2mp MBONs 15 min after training. (D) At 24 h CS- responses were significantly reduced over those of CS+ in β′2mp MBONs. (E) CS+ responses were relatively depressed over those of CS- in γ5β′2a MBONs 15 min after training. (F) At 24 h CS- responses were significantly reduced over those of CS+ in γ5β′2a MBONs. (G) CS+ responses were significantly facilitated over those of CS- in γ3,γ3β′1 MBONs 15 min after training. (H) CS+ responses were significantly depressed over those of CS- in γ3,γ3β′1 MBONs at 24 h. (I) CS+ responses were significantly reduced over those of CS- in γ2α′1 MBONs at 15 min. (J) CS+ responses became significantly potentiated over those of CS- in γ2α′1 MBONs by 24 h. (K) 15 min after training, CS- evoked a clear excitatory response over an obviously fluctuating baseline activity of γ1pedc>αβ MBONs, but no CS+ responses were evident. (L) By 24 h, no significant difference in CS+ versus CS- odor responses were evident in γ1pedc MBONs. MCH and OCT were interchanged as the CS+ and CS- odors in an equal number of trials Odor-evoked activity traces show mean responses (solid lines) ± SEM (shaded areas). Gray bars under the traces indicate 5 s odor stimulus. Quantifications represent normalized area under the curve (AUC) as mean ± SEM, with individual measurements shown as dots; paired data are connected by gray lines. Statistical comparisons were performed using Wilcoxon matched-pairs signed-rank test (n ≥ 7). Asterisks denote significance: *p < 0.05, **p < 0.01, ***p < 0.001, ****p < 0.0001.

The approach-directing γ2α′1 and γ1pedc>αβ MBONs also showed clear time-dependent changes in their odor-evoked responses. γ2α′1 MBONs responses to CS- were initially potentiated versus those of CS+ (consistent with prior data ^45^) but converted to greater CS+/CS- potentiation by 24 h. Odor evoked responses of γ1pedc>αβ MBONs at 15 min after training were less clear, likely because MBON activity is affected by nutrient-evoked oscillations of their cognate PPL1-γ1pedc DANs after sugar training. However, a strong relative depression of responses to the previously sugar-paired CS+ odor was evident, whereas by 24 h responses to the CS+ and CS- were equivalent and showed no obvious signs of plasticity. Therefore, γ1pedc>αβ MBON responses are indicative of malaise- directed interference with CS+ processing 15 min after training, which appears to be completed 24 h later.

### Fat body derived signals direct memory updating in the brain

Since amygdalin is metabolized in the gut, we used peptidomic analyses of sick fly hemolymph (Figure S11) to identify visceral signals that might mediate malaise-directed memory updating. We compared peptidomic profiles from flies trained with toxin-tainted sugar to those trained with sugar alone. 7 of the top 10 differentially abundant molecules (nothing reached statistical significance after multi-sample correction, but samples were notably variable) were known extracellular proteases and secreted proteins (Figure S12). Amongst these molecules, the Turandot protein TotA ^118–122^ and modular serine protease (modSP ^123^) have previously been implicated in the organismal response to toxic insult and are highly expressed in the fly’s fat body, an adipose and liver-like tissue that plays roles in detoxification, immune responses, and endocrine signaling. GCaMP7f expression in the head fatbody revealed higher baseline calcium levels 15 min after sucrose-amygdalin training, than in sucrose trained flies, consistent with tissue activation and release of hormonal signals (Figure 7A & 7B). We therefore tested whether fatbody production of Tot A (with Tot C as a control) and modSP was required for malaise-directed memory updating.

**Figure 7.**
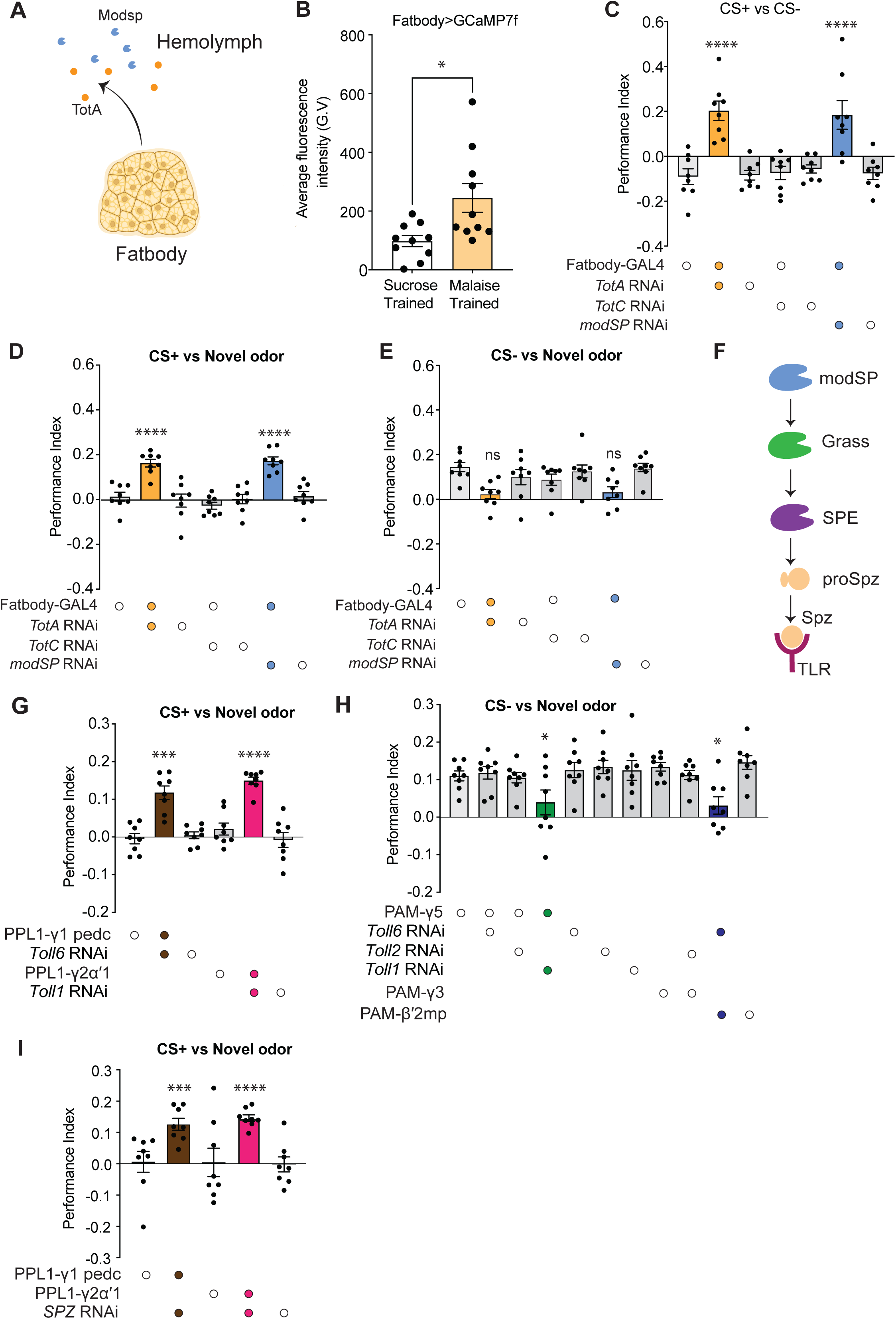
Fat body-derived signals direct memory updating in the brain and via autocrine Toll signaling in dopaminergic neurons. (A) Illustration of modSP and TotA release from fatbody in malaise trained flies. (B) Quantification of head fatbody intracellular calcium (Lpp-GAL4; GCaMP7f) measured 15 min after training in sucrose and malaise conditioned flies. (C) RNAi-mediated knockdown of *modSP*, *TotA*, but not *TotC*, in fatbody restores 24 h approach memory to malaise trained flies, when tested CS+ vs CS- odors. (D) RNAi knockdown of *modSP*, *TotA*, but not *TotC*, in fatbody restored 24 h CS+ approach memory when measured v novel odor. (E) RNAi knockdown of *modSP*, *TotA*, but not *TotC*, in fat body blocked formation of malaise-induced 24 h CS- approach memory measured v novel odor. (F) Schematic of proteolytic cascade leading to liberation of active Spätzle (Spz), the Toll receptor ligand. Autocatalytic modSP (modular Serine Protease) cleaves and activates the clip-protease Grass (Gastrulation Defective-Like Serine Protease), which in turn cleaves and activates SPE (Spätzle- Processing Enzyme). SPE cleaves pro-Spätzle (proSpz), generating Spz. (G) RNAi- mediated *Toll-6* knockdown in γ1pedc and *Toll-1 in* γ2α′1 PPL1 DANs restored 24 h CS+ approach memory measured v novel odor. (H) Knockdown of *Toll-6* in β’2mp and *Toll-1 in* γ5 PAM-DANs impaired malaise-induced 24 h CS- approach memory measured v novel odor. (I) Knockdown of *Spz* in γ1pedc or γ2α′1 PPL1 DANs restored 24 h CS+ approach memory measured v novel odor. Data in panel B were compared using Mann Whitney t-test, n=18, data points represent individual fly signals. Data in panel C were compared using one-way ANOVA with Tukey’s test. Data in panels D, E, G, H, I & J were compared using one sample t-test and Wilcoxon test, individual data points represent independent groups of approximately 200 flies. Asterisks denote significant differences: ^∗^*p* < 0.05, ^∗∗^*p* < 0.01, ^∗∗∗^*p* < 0.001, ^∗∗∗∗^*p* < 0.0001.

We used a fatbody specific GAL4 (apolipophorin, Lpp-GAL4 ^124^) to direct expression of transgenic RNAi targeting *modSP*, *TotA* or *TotC*. Although no effect was observed with *TotC* knockdown, fat body-restricted loss of either *modSP* or *TotA* abolished malaise-directed memory updating. 24 h CS+ v CS- memory performance was converted from relative CS+ avoidance back to CS+ attraction, thereby restoring sugar reward memory (Figure 7C).

When testing for the independent CS+ malaise memory, all controls and flies expressing *TotC* RNAi in fat body showed no 24 h CS+ approach, whereas those with *TotA* and *modSP* knockdown showed performance (Figure 7D), again consistent with re-emergence of sugar-rewarded CS+ approach memory. Similarly, while all control and *TotC* knock down flies showed clear 24 h CS- approach memory, no performance was evident in flies with fatbody *modSP* or *TotA* RNAi (Figure 7E). These *modSP* and *TotA* fatbody loss of function phenotypes therefore mimic those observed with disruption of consolidation using DPM neurons, and differential loss of 5-HT receptors in specific mushroom body innervating DANs.

### Memory updating requires autocrine Toll signaling in DANs

ModSP is a master autocatalytic protease that, in the context of gram-positive bacterial and fungal infection, can trigger a sequential cascade of Clip-proteases that ends with the Spaetzle-processing enzyme (SPE) cleaving and activating the cytokine and Toll-receptor ligand, Spätzle (Spz) ^124^ (illustrated in Figure 7F). Prior work has implicated Drosophila neurotrophin-2 (DNT-2) signaling through the Toll-6 receptor in Drosophila PPL1 and PAM dopaminergic neurons ^125^. We therefore queried our scSeq profiles of DANs for Toll-like receptors (TLRs) (Figure S13), which revealed differential expression of *Toll-1*, *Toll-2, Toll-6* and *Toll-7*. Most strikingly from the perspective of malaise memory, PPL1-γ1pedc and some PAM-β′2mp and PAM-γ5 DANs strongly expressed *Toll-6*, whereas PPL1-γ2α′1 most strongly expressed *Toll-1*. We also noted that γ3 DANs lack *Toll-6* but some weakly express *Toll-1* and *Toll-2*.

Led by expression profiles, we tested for roles of the relevant TLRs by driving DAN-specific expression of RNAi transgenes. Strikingly, RNAi knockdown of *Toll-6* in PPL1- γ1pedc and *Toll-1* in PPL1-γ2α′1 restored 24 h sugar reward CS+ memory to malaise-trained flies, but did not impair formation of CS- approach memory (Figure 7G & 7H). In contrast, removing *Toll-6* from PAM-β′2mp and *Toll-1* from PAM-γ5 DANs impaired formation of 24 h CS- approach memory (Figure 7I). Therefore, DAN subtype-restricted expression of specific TLRs is required for malaise-directed memory updating.

We hypothesized that modSP’s role in malaise memory updating could be to initiate the cleavage and activation of Spz to signal through Toll-6 and Toll-1 in the relevant DANs. Since our scSeq data revealed that PPL1- γ1pedc and PPL1-γ2α′1 DANs also express Spz- 1 (Figure S13), we tested whether the DANs themselves are the critical source of Spz.

Consistent with this idea, RNAi knockdown of *Spz* in PPL1-γ1pedc and PPL1-γ2α′1 DANs restored 24 h sugar reward memory to malaise-trained flies (Figure 7J). We therefore conclude that malaise memory updating is directed by modSP triggering autocrine Spz::Toll- 6 and Spz::Toll-1 receptor signalling in the PPL1-γ1pedc and PPL1-γ2α′1 DANs, which together block nutrient-dependent consolidation of the sugar reward memory. Autocrine Spz::Toll signalling in β′2mp and γ5 DANs could therefore also reverse relative CS+/CS- depression, giving rise to CS- approach memory.

### TotA is neuroprotective and may represent another malaise signal

Our TotA loss-of-function experiments produced similar results to that of loss of modSP. Prior work has suggested that circulating Turandot proteins protect tissues from the fly’s own immune response, and microbial attack, by binding to phosphatidyl serine (PS) when it is exposed on the outer leaflet of cell membranes of vulnerable cells ^118^. We therefore tested for a possible neuroprotective role of TotA during malaise-directed memory updating. We reasoned that TotA might be required to protect PPL1-γ1pedc DANs from elevated Toll receptor activation, which can result in PS exposure on the outer membrane. Prior work has established that PPL1- γ1pedc DANs convey the reinforcing effects of electric shock and also provide hunger-dependent control of appetitive memory expression. We therefore used these established roles to assay potential loss of PPL1-γ1pedc DAN function in malaise- trained *TotA* mutant flies (Figure 8). We first trained wild-type, *Tot*^AZ^ (these flies carry a deletion of the genomic cluster encoding TotA, TotB, TotC and TotZ) and *Tot*^AZX^ (AZ deletion and removal of TotX) mutant flies with the sugar/toxin malaise protocol, then 24 h later tested them for shock-reinforced aversive learning. As controls, parallel sets of flies of the same genotype were tested for aversive learning without preceding malaise training (Figure 8A). Wild-type flies showed similar aversive shock learning performance without and following malaise training (Figure 8B). However, aversive shock learning was significantly impaired in *Tot*^AZ^ and *Tot*^AZX^ flies following malaise training (Figure 8C & D).

**Figure 8.**
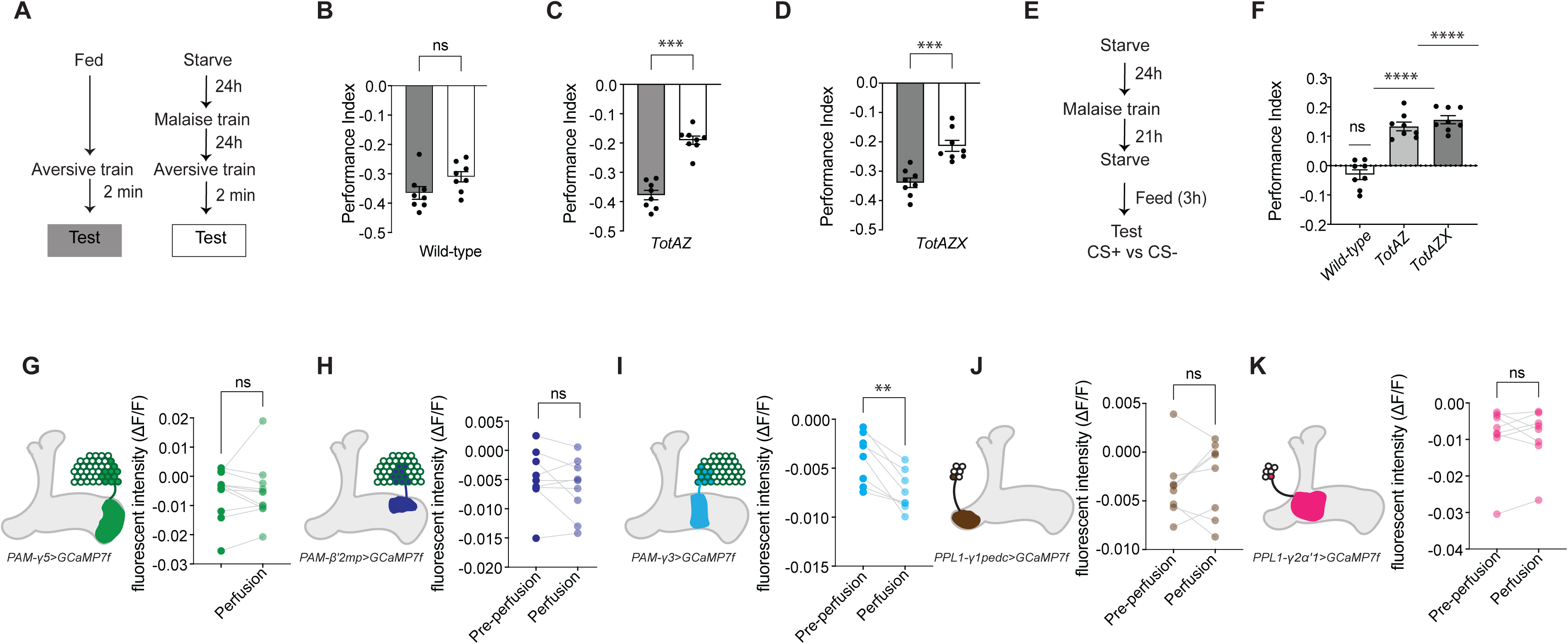
TotA provides DAN resilience and may also alter γ3 DAN activity. (A-D) Tot genes maintain PPL1- γ1pedc DAN activity following malaise learning. (A) Experimental protocol to assess maintenance of aversive learning following malaise training. (B). Wild-type flies show no difference in their shock-reinforced aversive learning ability 24 h following malaise conditioning. (C & D) *Tot*^AZ^ (C) and *Tot*^AZX^ (D) mutant flies exhibit defective aversive shock learning 24 h after malaise training. (E) Experimental protocol to assess maintenance of state-dependent motivational role of PPL1-γ1pedc DANs after malaise memory. (F) wild-type flies show no 24 h memory performance when *ad libitum* fed for 3 h prior to CS+ v CS- memory testing. In contrast, *Tot*^AZ^ and *Tot*^AZX^ mutant flies express CS+ approach memory, that is resistant to suppression by feeding. Panels (B, C, D & F), individual data points represent independent groups of approximately 200 flies. (G-K) Imaging DAN responses to TotA application. Left: schematics of γ5, β’2mp, γ3, γ2α′1 & γ1pedc DANs. Right, Calcium imaging values (DANs express UAS-GCaMP7f) before and during TotA peptide application (in presence of TTX). Only PAM-γ3 DANs show a consistent and significant decrease in their intracellular calcium to TotA. Data in panels G-K, individual data points show each fly’s response. Data in panels B-D and F were compared using Mann Whitney paired t-test, n ≥ 8. Data in Figure (I) were compared using one sample t and Wilcoxon test. Data in panels G and K were compared using t tests (non-parametric tests) with Wilcoxon matched pairs signed rank test, n≥8. Asterisks denote significant differences: ^∗^*p* < 0.05, ^∗∗^*p* < 0.01, ^∗∗∗^*p* < 0.001, ^∗∗∗∗^*p* < 0.0001.

We next trained wild-type, *Tot*^AZ^ and *Tot*^AZX^ flies with the malaise protocol then tested whether the expression of the restored 24 h sugar rewarded memory could be suppressed by feeding the flies ad libitum for 3 h prior to memory testing. Whereas performance of wild-type flies was abolished in food satiated flies, expression remained in *Tot*^AZ^ and *Tot*^AZX^ flies (Figure 8F). Therefore, both aversive learning and hunger-dependent control of appetitive memory expression are defective in *Tot* mutant flies following malaise training, consistent with loss of a neuroprotective function for TotA.

We also considered whether TotA could alter activity of malaise-relevant DANs. Strikingly, bath application of a putative active fragment of TotA (RLYPSLTPEERESIDKF) to dissected brains expressing DAN-specific GCaMP7f, revealed a strong and consistent reduction of intracellular calcium only in the PAM-γ3 DANs (Figure 8I). Since the PAM-γ3 DANs were the only memory-updating group that did not show obvious consequences of TLR loss of function, we speculate that TotA may also have a novel and unexpected signalling role in these DANs.

## Discussion

Some toxins in food are perceived as tasting bitter and are therefore innately avoided prior to ingestion. For those that bypass this peripheral barrier of food selection, and that do not kill, post-ingestive effects allow animals to learn to avoid toxin-associated food in future. Our understanding of the mechanisms that assign malaise value to feeding memories is rudimentary.

We demonstrated that *Drosophila* learn to associate malaise with odors experienced just before, and during, eating toxin-tainted sucrose. Immediately after training, flies show preference for the sucrose/toxin associated odor but within 30 min preference is nullified, and by 6 h they express preference for the odor presented prior to feeding. Flies modify their initial responses to both odors after intoxication. Their immediate approach memory for the previously sugar/toxin associated odor is relatively quickly erased while approach to the non-feeding paired odor emerges more slowly. These observations are highly reminiscent of the dynamic olfactory preferences of *Caenorhabiditis elegans* nematodes fed with pathogenic and nonpathogenic bacteria, which also have approach and aversion components ^18^.

### Consolidation released 5-HT allows integration of post-ingestive information

Finding that malaise-conditioned behavior switched valence around 15-45 min after training led us to hypothesize that post-ingestive malaise information was integrated while memories are consolidated. Prior work established that the serotonergic DPM neurons are required at a similar time after training to consolidate nutrient-dependent sucrose-reinforced LTM ^27, 68^. Moreover, a recent study showed that DPM released 5-HT defines the effective ISI for odor shock pairing ^11^. Artificially stimulating DPM neurons, increasing their synthesis of 5-HT, or impeding 5-HT uptake all extended the ISI from 17 to >40 s. Our work here suggests that DPM neuron released 5-HT can extend the effective ISI much further after a feeding event into the consolidation window so that post-ingestive reinforcement can be integrated.

We found that DPM block 15-30 min after training abolished formation of malaise memories but still permitted nutrient-dependent consolidation of sucrose LTM and that this distinction could also be observed with loss of specific 5-HT receptors from DANs that are involved in both processes. This suggests that these two post-ingestive signals are integrated during temporally separable periods within the consolidation window, with amygdalin coming first. As discussed below, this relative timing aligns with our mechanistic explanation for how malaise value is assigned to both the feeding-event odors. Since the metabolization and detection of beneficial or harmful effects of different substances are likely to have discrete dynamics, there is utility to consolidation spanning a window of time large enough to accommodate a variety of signals. It will be interesting to investigate how post-ingestive effects of other nutrients, toxins and pathogens are learned.

Finding that 5-HT directed modulation of DANs is required to permit integration of malaise information extends the proposed roles for this modulator in malaise learning. Both insects and mammals have gut-associated cells that produce and can release 5-HT into the hemolymph/ blood stream on feeding ^80–83^. 5-HT from enterochromaffin cells in the mammalian intestine signal nausea and induce vomiting via activating 5-HT3 receptors in the vagus nerve ^84^. Other studies in *Drosophila* have reported roles for specific 5-HT neurons in regulating food-seeking and consummatory behavior ^85–87^. More broadly, 5-HT has been frequently associated with inhibitory and aversive behaviors and in opposition to the action of dopamine ^88–91^. However, it is important to emphasize that DPM neuron released 5-HT is required to consolidate memories, irrespective of their valence ^27, 66, 67^.

Nevertheless, it will be important to understand the full extent of 5-HT regulated processes in malaise learning, and whether different sources of release engage common targets and mechanisms.

### Humoral modSP represents malaise through autocrine TLR signaling in DANs

Finding modSP to be upregulated in the hemolymph of intoxicated flies, pointed to the possible involvement of Toll signaling in memory updating. Surprisingly, we discovered that key PPL1-γ1pedc and PPL1-γ2α’1 DANs co-express Spz and TLRs giving them the potential to be regulated by autocrine Toll signaling. Since secreted Spz needs to be cleaved to be active, humoral modSP can direct DAN-specific TLR signaling by triggering the clip-protease cascade that culminates in local Spz cleavage. These DANs are therefore effectively primed to signal malaise, but their DAN-released Spz is cryptic until cleaved.

ModSP has been most studied for its role in the fly’s Toll-dependent immune response to fungal and gram-positive bacterial infection. Amygdalin is abundant in the seeds of many plants, especially those of the Roseacea family such as almonds (where it derives its name from the Greek, amygdalē), apricots, peaches and cherries. Damage to the flesh of these fruits releases enzymes that hydrolyze amygdalin into hydrogen cyanide and benzaldehyde to deter consumption. In our experiments we expect amygdalin to be hydrolyzed after ingestion by β-glucosidase in the fly’s gut ^100^ (co-feeding amygdalin and glucosidase inhibitor blocks malaise learning, Figure S8). Given the reactivity of hydrogen cyanide, we expect tissue damage to be a trigger of fatbody release of modSP and TotA.

Interestingly, prior studies of CTA and TPOA in the fly relied on post-ingestive detection of the virulent intestinal pathogen *Pseudomonas entomophila* ^19, 20^. This bacterium produces monalysin, a pore-forming toxin that also damages the gut ^101^. In addition, preference for harmless over harmful pathogen was expressed about 15 min after first ingestion ^95^. Since a broad range of pathogens activate Spz by intervening with specific clip-proteases in the sequential modSP regulated cascade, we speculate that many other pathogens and toxins will ultimately signal post-ingestive aversive reinforcement via TLR signaling in the same DANs identified here. It is notable that Kobler et al (2020) ^19^ did not observe a major contribution of DANs in their CTA assay and instead concluded a role for immune signal detection in octopaminergic neurons. Our prior work suggests that OA is critical for sweet taste reinforced olfactory STM, although we believe OA acts through DANs ^32, 40^. Given the importance of DANs for many forms of olfactory learning, it could be that they are more critical for TPOA than CTA.

It will be important to understand the physiological consequences of TLR signaling in the respective DANs. Although we found DAN-specific roles for different TLRs, the resulting effects of their modulation was consistently to reverse the sugar-reward evoked plasticity of relative CS+/CS- odor responses observed in their cognate MBONs. We note that the malaise-updating PPL1-γ1pedc and PAM-γ5 DANs have been reported to co-release nitric oxide (NO) and that NO-induced behavioral plasticity was found antagonize the role of dopamine ^127^. Since our DAN scSeq profiles show that NOS expression is restricted to these and the other malaise-relevant PPL1- γ2α’1, PAM-γ3, PAM-γ5 and PAM-β′2mp DANs, we therefore predict that malaise-directed memory updating is the physiologically relevant role for NO release from DANs. This notion is further supported by the knowledge that TLR activation can trigger NF-kB (Dorsal/Dif-A)-dependent expression of immune response genes ^128,129^, that includes inducible nitric oxide synthase (iNOS) in mammalian macrophages ^130,131^. Moreover, shorter non-nuclear Dorsal-B and Dif-B isoforms have been reported to have a synaptic function in the fly ^132,133^. It therefore seems possible that nuclear and synaptic TLR signaling could function together to switch DAN transmitter usage to favor NO over DA, to direct memory updating.

### Credit assignment

Studies of conditioned flavor preference established that differential taste is required for animals to learn associations between distal cues such as odors, colors and objects with post-ingestive nutritive reward ^3^. In our assay the second of the two odors needs to be presented with sweet sugar and toxin. No odor preference was developed if the first of two odors was paired with sugar and toxin. We assume that in this case the flies are unable to disambiguate the two odors because the slow processing of toxin ingested during the first odor presentation contaminates the second odor experience. We also observed no conditioned odor preference if flies were trained with one odor presented with sucrose and the other with sucrose plus toxin, consistent with a requirement for differential taste and the need to distinguish between the feeding-relevance of the two odors. We found that malaise forms parallel memories for both odors experienced during training and that odor-sugar learning establishes labile traces for each odor in the KC-MBON network that post-ingestive nutritive or malaise signals later act upon.

We found that removing inhibitory 5-HT1A and Spz::Toll-6 from PPL1-γ1pedc DANs abolished the assignment of malaise to the CS+ odor and instead produced re-emergence of 24 h CS+ approach memory. Moreover, directly blocking PPL1-γ1pedc DANs 15-45 min after training also recovered CS+ approach. Interestingly, prior work has shown that the oscillatory activity of PPL1-γ1pedc DANs is increased 30 min after training with nutritious D-glucose, but not with non-metabolizable L-glucose, and that their activity during this period is critical for consolidation of these STMs into nutrient-dependent LTM ^33^. Our functional imaging of the cognate γ1pedc>αβ MBON 24 h after malaise training revealed no evidence for plasticity of relative odor responses. We therefore conclude that malaise signaling to PPL1-γ1pedc DANs erases sugar-rewarded CS+ approach memory by interfering with its nutrient-dependent consolidation.

We also found that the inhibitory 5-HT1B receptor and Spz::Toll-1 were required in PPL1- γ2α’1 DANs for malaise directed loss of CS+ approach memory. Consistent with prior work after sucrose training ^45^, we initially observed potentiated responses to the CS- odor in the γ2α’1 MBONs. Given that γ2α’1MBONs are approach-directing ^92^, CS- potentiation is effectively a ‘latent’ CS- approach memory. Moreover, another study demonstrated that PPL1-γ2α’1 DAN activity after training can reverse learned contingencies measured in the γ2α’1 MBON ^93^, which provides a precedent for all of the malaise-directed reversals we observed here. In addition, a PPL1-γ2α’1 DAN directed reversal is consistent with our finding that stimulation of these DANs 15-45 min after training restored CS+ approach, because PPL1-γ2α’1 DAN activity would be predicted to turn relative CS- potentiation of responses in γ2a’1 MBONs into CS+ potentiation.

Development of 24 h CS- approach required excitatory 5-HT7 receptors in PAM-γ3, 5-HT7 and Toll-6 in PAM-β’2mp, and inhibitory 5-HT1A and Toll-1 in PAM-γ5 DANs. Moreover, whereas PAM-γ3 and PAM-β’2mp DAN activity was required 5-45 min after training for CS-memory, PAM-γ5 stimulation at this time limited it. We previously implicated PAM-γ3 and PAM-β’2mp DANs in formation of a slowly emerging CS- safety memory following multiple spaced trials of aversive odor-shock conditioning ^47^. Although malaise learning occurs after a single learning trial and the potential assignment of CS- safety occurs after the odor is presented, the neural processes and the dynamic of each CS- memory appear remarkably similar. Learning that an odor is not accompanied by shock punishment also required PAM- γ3 DANs and a resulting loss of feedforward inhibition from the γ3β’1 MBONs was proposed to release PAM-β’2mp DANs to write the CS- safety memory ^47^. Our functional imaging revealed that the relative odor responses of γ3β’1 MBONs were significantly reversed 2 h after malaise training, and before that of the β′2mp and γ5 MBONs, consistent with their plasticity being key for development of CS- approach memory at these other junctions. We propose that TLR-modulated DAN activity is sufficient to reverse the relative odor-evoked KC-MBON plasticity formed by sugar reward. Although more work will be required, we speculate that PAM-γ3 DANs might also be modulated by TotA.

We noted that 6 h after training the flies show a stronger approach to the CS- odor when tested versus a novel odor rather than the CS+. We therefore speculate that the flies also learn to generalize their aversion to novel odors for a few hours after experiencing malaise. We are currently unsure how such generalized odor aversion would manifest in the fly brain.

Finding that malaise learning interferes with consolidation of prior odor-sugar memory provides a logic for why post-ingestive reinforcement can sometimes be ambiguously assigned to several prior feeding experiences. We predict that other experiences that occur before a memory of a prior event is consolidated will also be assigned a similar malaise value. However, feeding events that occur after the toxin-tainted experience has been consolidated should be treated as a separate and distinguishable event. With this reasoning, the time to completion of consolidation represents an event boundary.

There are many remaining questions regarding formation of malaise memories. For example, it is unclear how the system can accurately assign the malaise updates to the correct odor representations, while reversing the relationship between the two odor memories at each KC-MBON junction during memory consolidation. We propose that the process involves spontaneous KC activity after training. α’β’ KC output is required at a similar time to that of DPM neurons to consolidate sugar memory ^26, 27^, KCs are required to drive 5-HT release from DPM neurons ^11^, and the key malaise-memory directing PPL1- γ2α’1, PAM-γ3 and PAM-β’2mp DANs all receive direct and second order MBON feedback input from these KCs ^71^. In addition, γ2α’1 MBONs provide feedforward excitation to the PAM-β’2mp reward DANs ^94^, which could potentially allow spontaneous α’β’ KC activity to replay the prior ‘latent’ learned potentiation of CS- connections onto γ2α’1 MBONs to excite PAM-β’2mp DANs to form the emerging CS- approach memory at that KC-MBON connection. Moreover, γ3β’1 MBONs provide feedforward inhibition onto PAM-β’2mp DANs and they initially harbor relative CS+/CS- odor responses that are reversed to that of γ2α’1 MBONs. Therefore, α’β’ KC activity after training can reactivate MBON network representations of both CS+ and CS- memories. Interestingly, a recent study reported that post-ingestive malaise reactivated novel flavor representations in neurons in the mouse amygdala that were previously observed during a recent meal ^95^. Such a malaise-driven reactivation of odor/taste representations in the flies’ KCs would aid our observed memory updating.

### Parallel memory representations

Several of our previous studies have demonstrated that fly behavior is informed by complementary or competing parallel memories ^45–47, 98, 99^. Most relevant here, following training with bitter-tainted sucrose flies, show a short-lived aversive memory that later converts to approach driven by a parallel sugar-rewarded appetitive memory ^98^. In addition, 24 h after multiple spaced trials of aversive conditioning fly behavior is guided by an avoidance memory for the shock paired odor and a safety memory for the explicitly unpaired odor ^47^. We found that malaise learning also forms parallel memories for both odors, but 24 h after training behavior appears to be solely directed by a safety memory for the non-food paired odor – no CS+ approach or avoidance was evident at this time.

We were surprised to find that malaise abolished CS+ approach and did not redirect behavior towards CS+ avoidance. Investigations of memory extinction in *Drosophila* established that the omission of expected reward or punishment (prediction errors) drive the formation of competing parallel memories of opposite valence, that nullify or extinguish conditioned behavior ^45, 46, 77^. By remembering reinforced and non-reinforced trials the fly can keep track of the reliability of sequential experience. However, our findings with 5-HT1A in the aversively reinforcing PPL1-γ1pedc DANs indicate that the fast malaise-directed loss of conditioned approach to the previously rewarded CS+ does not involve formation of a parallel competing aversive CS+ malaise memory. Instead, our studies suggest that malaise interferes with consolidation of the earlier sugar-rewarded short-term odor memory. To our knowledge this is a unique example of new information leading to erasure of a prior memory, rather than retention of competing evidence. Perhaps the potential lethality of amygdalin consumption makes it too risky to retain a reward memory that might lead to the fly ingesting it again. It will be interesting to further investigate the nature and persistence of this learned aversion.

### Is post-ingestive learning the purpose of memory consolidation?

The purpose of consolidation has been debated, from simply representing a constraint imposed by biological hardware, to a period after training during which new memories are malleable and can be readily linked to other information to produce more meaningful memories/ narratives ^102–105^. Our data support the latter perspective. Moreover, since olfaction, gustation and general visceral afference are considered to be special senses because they are directly conveyed to the limbic system ^106^, it seems plausible that integration of post-ingestive information into prior feeding-related memories is an ancestral purpose of consolidation.

## Acknowledgements

We thank G. Wright for inspiring this work, T. Behrens and members of the Waddell group for discussion. We thank R. Brain, R.Busby and F. Woods for technical assistance, A. Mishra for help with hemolymph collection, and the Goodwin lab for locomotor tracking. We are grateful to B. Lemaitre, the Bloomington Drosophila Stock Center (NIH P40OD018537) and the Vienna Drosophila Resource Center for fly stocks. S.W. was funded by a Wellcome Principal Research Fellowship (200846), a Wellcome Discovery Award (225192), an ERC Advanced Grant (789274), and Wellcome Collaborative Awards (203261 and 209235).

## Author contributions

Designed research B.S., C.T., S.W., Performed research B.S., C.T., Analyzed data B.S., C.T., S.W., Resources S.W., Writing S.W. B.S., C.T., Supervision S.W., Funding Acquisition S.W.

## Declaration of interests

The authors declare no competing interests.

---

**Figure S1, related to Figure 1. Locomotion is temporarily reduced following malaise training.**

(A) Upper panels, screenshot of flies in the 20mm diameter arena. Lower panels, representative locomotor traces of sugar trained (blue) and malaise (red) trained flies. (B) Quantification of average velocity per fly (cm/s). Columns 1 and 2; malaise-trained flies showed significantly reduced locomotion compared with sugar-trained flies at 15 min. Columns 3 and 4; malaise-trained flies show partial recovery of locomotion after 24 h. Data analyzed using one-way ANOVA with Tukey’s post hoc test (n ≥ 16). Each point represents an individual fly. Data presented as mean ± SEM. Significance indicated as: *p < 0.05, **p < 0.01, ***p < 0.001, ****p < 0.0001.

**Figure S2, related to Figure 1. Optimization of malaise learning.**

(A) 2 min and 24 h memory performance of starved flies trained with 0.0, 0.2, 0.4, 0.6, 0.8 and 1.0 % concentrations of amygdalin in sucrose. 0.6 % produced optimal performance. (B) 1. Flies trained using our chosen malaise protocol, odor 1 with nothing then odor 2 with sucrose/amygdalin, show significant 24 h CS+ v CS- odor preference. 2. Flies trained with the first of the two odors paired with sucrose and 0.6 % amygdalin do not show significant 24 h odor preference. 3. Flies do not show significant 24 h odor preference if differentially conditioned by pairing one odor with sucrose and the other with sucrose plus 0.6 % amygdalin. (C) 2 min and 24 h memory performance of starved flies trained with 0.2, 0.4, 0.6, and 0.8% of quinine in sucrose. (D) Odor acuity (preference between odor v air) measured 24 h after feeding either sucrose or sucrose plus amygdalin. (E) Malaise training does not significantly alter sucrose consumption measured 24 h after training. Data in panels A and C were compared using one-way ANOVA with Tukey’s test, n = 4-8. Data in panels B, D and E were compared using paired t-tests and Wilcoxon tests, n ≥8; Data are mean ± SEM, asterisks denote significant differences: ^∗^*p* < 0.05, ^∗∗^*p* < 0.01, ^∗∗∗^*p* < 0.001, ^∗∗∗∗^*p* < 0.0001.

**Figure S3, related to Figure 3. Conditioned behavior changes between 15 and 45 min after malaise training.**

A magnified view of memory performance timelines during the first 2 h after training, measuring CS+ v CS-, CS+ v Novel odor, and CS- v Novel odor. Both the intoxication- induced decrease in CS+ approach and emergence of CS- approach occur between 15 and 45 min after training. Asterisks denote significant differences: ^∗^*p* < 0.05, ^∗∗^*p* < 0.01, ^∗∗∗^*p* < 0.001, ^∗∗∗∗^*p* < 0.0001.

**Figure S4, related to Figure 4. Serotonin receptors are differentially expressed in PAM and PPL1 DANs.**

Dot plot showing normalised, unscaled average expression levels of 5-HT receptor mRNA in DANs. Only average expression levels present in more than 40% of cells in each cluster are shown. DANs are ordered and grouped based on the identity of the KC type and mushroom body compartments that they innervate.

**Figure S5, related to Figure 4. Malaise-induced CS- approach memory requires 5-HT receptors in rewarding DANs.**

Left, RNAi-mediated knock down of 5-HT1A, 5-HT1B, 5-HT2A, 5-HT2B and 5-HT7 in R58E02-GAL4 expressing PAM DANs. Data was compared by using one-sample t and Wilcoxon test, *n* ≥8; Data are mean ± SEM; dots are individual data points. Right, Diagram summarizing differential 5-HT receptor coupling.

**Figure S6, related to Figure 4. Differential 5-HT modulation of DAN subtypes is required for CS- malaise memory.**

Detail of an initial screen measuring 24 h CS- approach memory in flies with specific 5-HT receptors knocked down with RNAi in specific DAN subtypes. Data in panel A was compared using one-way ANOVA with Tukey’s test, *n* ≥8; Data are mean ± SEM; dots are individual data points, which represent independent groups of approximately 200 flies. Asterisks denote significant differences: ^∗^*p* < 0.05, ^∗∗^*p* < 0.01, ^∗∗∗^*p* < 0.001, ^∗∗∗∗^*p* < 0.0001.

**Figure S7, related to Figure 5. PAM-γ3 DAN manipulation after training also impairs sucrose memory.**

(A) Behavior protocol used in panels B and C. (B) A 30 min UAS-*Shi^ts^* mediated block of γ3 DANs from 15 min after training impairs sucrose memory (CS+ vs CS-). (C) 30 min UAS- *dTrpA1* mediated activation of γ3 DANs from 15 min after training impairs sucrose memory (CS+ vs CS-). Data in panels B and C were compared using one-sample t-test and Wilcoxon test, *n* ≥12 (Figure B), *n* ≥8 (Figure C). Data are mean ± SEM; dots are individual data points, which represent independent groups of approximately 200 flies. Asterisks denote significant differences: ^∗^*p* < 0.05, ^∗∗^*p* < 0.01, ^∗∗∗^*p* < 0.001, ^∗∗∗∗^*p* < 0.0001.

**Figure S8, related to Figure 1. Flies prefed with β-glucosidase inhibitor exhibit mild restoration of 24 h CS+ approach in malaise trained flies.**

(A) Behavioral paradigm. (B) Flies fed with the irreversible inhibitor of β-glucosidase conduritol B epoxide before training show a significant difference of 24 h memory performance compared to those not fed inhibitor. Data were compared using Mann Whitney paired t-test, n ≥ 8. Data are mean ± SEM; dots are individual data points which represent independent groups of approximately 200 flies. Asterisks denote significant differences: ^∗^*p* < 0.05, ^∗∗^*p* < 0.01, ^∗∗∗^*p* < 0.001, ^∗∗∗∗^*p* < 0.0001.

**Figure S9, related to Figure 6, MBONs required for malaise memory.**

(A and B) Upper panels, temperature shifting protocols imposed on training and testing timelines. (A) UAS-*Shi^ts^* mediated block of γ3,γ3β’1 and γ5β’2a/β’2a MBONs for 30 min before and during testing impairs expression of malaise-induced CS- approach memory measured at 24 h v novel odor. (B) UAS-*Shi^ts^* mediated block of γ2β’1 and γ1pedc>αβ MBONs for 30 min before and during testing does not reveal a difference for malaise- induced CS+ avoidance memory measured at 24 h v novel odor training. Note there should be no performance in the controls. (C) Illustration of the MBON compartments that mediate malaise memories. (D) Malaise MBONs characterized by their primary neurotransmitters and the valence of their activation. The observed plasticity of the odor responses of these MBONs largely explains the behaviour observed at 24 h after training. Data in panels A and B were compared using one-way ANOVA with Tukey’s test, *n*=8; Data are mean ± SEM. Individual data points represent independent groups of approximately 200 flies. Asterisks denote significant differences: ^∗^*p* < 0.05, ^∗∗^*p* < 0.01, ^∗∗∗^*p* < 0.001, ^∗∗∗∗^*p* < 0.0001.

**Figure S10, related to Figure 6. KC-MBON plasticity measured 2 h after malaise training.**

(A) Protocol to measure odor-evoked responses in MBONs at 2 h after training. (B-F) Left panels, anatomy and imaging planes of MBONs: β’2mp, γ5B’2a, γ3,γ3β’1, γ2α’1 & γ1pedc>αβ; middle panels - odor-evoked activity traces show means (solid line) with SEM (shadow). Gray line underneath indicates 5 s odor exposure. Right panels - bar graphs display normalized area under the curve as means ± SEM. Individual data points are displayed as dots, and paired measurements are connected by gray lines. (B) No differences are apparent between CS+ and CS- odor evoked responses of β’2mp MBONs at 2 h. (C) No differences are apparent between CS+ and CS- odor evoked responses of γ5β’2a MBONs at 2 h. (D) At 2 h after malaise training, the relative responses of CS+ v CS- odors have already reversed in γ3,γ3β’1 MBONs, from those observed at 15 min (Figure 6G). (E) No differences are apparent between CS+ and CS- odor evoked responses of γ2αγ1 MBONs at 2 h. (F) CS- odor evoked responses are significantly reduced to those of CS+ responses in γ1pedc>αβ MBONs at 2 h. CS+ data correspond to average of experiments in which 50% of trials used MCH as CS+ and 50% used OCT. Same applies for CS- data. Asterisks denote significant difference between average responses to CS+ and CS−. Figures (B-F) were compared by using Wilcoxon matched -pairs signed rank test, n≥8. Asterisks denote significant differences: ^∗^*p* < 0.05, ^∗∗^*p* < 0.01, ^∗∗∗^*p* < 0.001, ^∗∗∗∗^*p* < 0.0001.

**Figure S11, related to Figure 7. The hemolymph collection for proteomics study**

(A) Protocol to train flies before hemolymph collection. (B) Steps described in detail for hemolymph extraction. (C) Steps followed for commercial proteomics of hemolymph.

**Figure S12, related to Figure 7. Top 10 candidate differentially detected hemolymoh peptides between sucrose-toxin and sucrose trained flies.**

(A) Heatmap representation of protein expression. Values were scaled per protein (row), allowing comparison of expression changes between experimental groups. The top 10 candidates are highlighted and annotated with their UniProt ID and refer to: Q9VEY0 - CG8927; A0A1B2AJ59 – GEO10156p1; A7DYW5 – CG17012; Q8IN44 – TotA; Q9VER6 – modSP; Q7K7G5 – EG:125H10.1 protein; F3YDP5 – CG43074-RA; X2JCV2 – LSP2; Q9VW34 – SERP; Q0GT46 – CG9897.

**Figure S13, related to Figure 7. Differential Toll receptor and Spz expression in PAM and PPL1 DANs.**

Dot plot showing normalized, unscaled average expression levels of Toll receptor mRNA in DANs. Only average expression levels present in more than 40% of cells in each cluster are shown.

## Methods

### Fly husbandry

All *Drosophila melanogaster* strains were maintained at 25 °C and 60 % humidity under a 12:12 h light:dark cycle (lights on from 08:00 to 20:00). For all behavioral assays, flies were reared on standard yellow cornmeal agar food consisting of deionized water, 7.2 g l^−1^ agar (Fisher Scientific), 25 g l^−1^ autolyzed yeast extract (Brian Drewitt), 47.3 g l^−1^cornmeal (Brian Drewitt), 100 g l^−1^ dextrose (D-glucose anhydrous, Fisher Scientific), 2.2 g l^−1^ Tegosept (methyl 4-hydroxybenzoate) (Sigma-Aldrich), and 8.4 ml l^−1^ ethanol (Sigma-Aldrich). Food was prepared by boiling. For optogenetic experiments, adult flies were reared in complete darkness for 3 days prior to deprivation on the same yellow cornmeal agar food supplemented with 1 mM all-*trans*-retinal (Sigma-Aldrich). For all physiological experiments, flies eclosed on brown agar food containing deionized water, 6.75 g l^−1^ agar, 25 g l^−1^ yeast, 62.5 g l^−1^ cornmeal, 37.5 ml l^−1^ molasses (Brian Drewitt), 4.2 ml l^−1^ propionic acid (Fisher Scientific), 1.4 g l^−1^ Tegosept and 7 ml l^−1^ ethanol. 7-to-9-day old adult flies were used for all experiments, and they were transferred to fresh vials a day before experimentation.

### Fly strains and genetic crosses

The control wild-type *Canton-S* strain originated in Chip Quinn’s lab (Massachusetts Institute of Technology, Cambridge, MA, USA). A variety of GAL4 lines were employed for targeted gene expression; *TH*-GAL4 ^107^, *R58E02*-GAL4 ^38^, and *VT64246*-GAL4 ^67^. To achieve more refined expression patterns, we used a panel of split-GAL4 lines produced at HHMI Janelia Research Campus ^108^: MB630B, MB296B, MB058B, MB312C, MB087C, MB320C, MB109B, MB315C, MB441B, MB213B, MB043C, and MB056B. Expression patterns can be viewed at http://splitgal4.janelia.org/cgi-bin/splitgal4.cgi. Optogenetic experiments involved expressing UAS-*GtACR1*^61^ under the control of specific GAL4 drivers. For thermogenetic activation and inhibition UAS-*dTrpA1* ^79^ and UAS*-Shi^ts^*^1^ ^65^ were expressed under control of specific GAL4 drivers. RNAi-mediated knockdown of serotonin receptors was performed by crossing GAL4 driver flies to UAS-*5-HT2A RNAi* (VDRC #102105), UAS-*5-HT2B RNAi* (BDSC #60488), UAS-*5-HT7 RNAi* (BDSC #27273), UAS-*5-HT1A RNAi* (VDRC #106094), and UAS-*5-HT1B RNAi* (BDSC #33418) lines ^109^ ^110^. For two-photon calcium imaging, the 20XUAS-*GCaMP7f* ^111^ reporter was expressed using appropriate GAL4 lines. Male and female flies were used in all behavioral and imaging assays.

### Naïve choice assay

To assess taste preference, 3–5 days old male and female flies (approximately 100 per trial) were starved for 20-24 h in vials containing 1% agar and a moistened filter paper.

Subsequently, flies were placed at the choice point of a T-maze. Two distinct odorized agar solutions were presented at the ends of the arms for a 2 min: (A) 1 M sucrose in 1% agar versus (B) 1 M sucrose + 0.6% quinine in 1% agar, or (A) 1 M sucrose in 1% agar versus (B) 1 M sucrose + amygdalin in 1% agar.

The attraction index was quantified as:

N (Sucrose + Toxin)-N(Sucrose)/N(Total)

Where N(Total) is the total number of flies tested.

### Proboscis extension reflex assay

3- to 5-day-old wild-type flies were food-deprived for 20-24 h. Flies were anesthetized for 1 min by placing them in a cold test tube immersed in a 4 °C ice bath, then mounted dorsally onto glass slides using a drop of nail polish and allowed to recover for 1.5 h at 25 °C and 60 % relative humidity. PER was elicited by sequentially stimulating either the foreleg or labellum with a rolled Kimwipe wick soaked in water (negative control), the test compound, and 3 M sucrose (positive control). Each test compound was presented three times per fly, with each fly receiving only one test substance flanked by water and sucrose controls. PER means response was quantified as the percentage of presentations eliciting proboscis extension. Flies exhibiting proboscis extension to water or failing to respond to 3 M sucrose were excluded from the analysis.

### Ingestion assay

All flies were food deprived before testing for 16–20 h in 1% agar vials containing a piece of filter paper. Sucrose was prepared as 1 M solutions with 0.4 % FD&C Blue No. 1 food dye (Spectrum Chemical) with 1% agar. 5 ml was pipetted onto an 8 X 5 cm piece of paper and allowed to settle. Papers were inserted in the training chamber of the olfactory conditioning apparatus. Around 80 flies were given 2 min to feed in the presence of airflow. Flies were then removed from the training chamber and immediately frozen at −20 °C to prevent excretion. Twenty flies were homogenized in 500 μl phosphate-buffered saline (1.86 mM NaH2PO4, 8.41 mM Na2HPO4, and 175 mM NaCl) and centrifuged at 14000 rpm for 3 min to clear debris. Supernatant was diluted with 100 μl PBS and centrifuged again at 14000 rpm for 3 min. Dye in the supernatant was quantified by measuring absorbance at 625 nm using a NanoDrop. Test compounds; sucrose + quinine (1M sucrose + 0.6% quinine and 1% agar solution with 0.4% FD&C Blue No. 1 food dye) and sucrose+ 0.6% of amygdalin (1M sucrose with 1% agar solution with 0.4% FD&C Blue No. 1 food dye).

### Gut dissection and food passage visualization

Following a 2 min feed on either 1 M sucrose solution (containing 0.4% FD&C Blue No. 1) or 1 M sucrose + 0.6% of amygdalin (containing 0.4% FD&C Blue No. 1), flies were rapidly chilled on ice at 4 °C. Subsequently, the entire digestive tract was dissected under a microscope. The presence and extent of blue food dye within the gut was visually assessed using a microscope to determine the progression of food through the digestive system.

### Malaise conditioning

Mixed sex populations of 3- to 5-day-old flies raised at 25 °C were tested together in all behavior experiments. Prior to training, groups of ∼100 flies were food-deprived for 20-24 h in vials containing 2-3 ml 1% agar and a strip of filter paper. For malaise CS+, a mixture of 1M sucrose, 0.6% of amygdalin or quinine along with 1% molten agar prepared. The solution spread in an even layer on piece of filter paper, backed with parafilm. For sucrose CS+ (1M sucrose along with 1% molten agar) was prepared. After drying for 3h, the papers were rolled into T-maze training tubes. For CS-, water-soaked dry filter paper used. Flies were trained by first exposing them to one odor (the CS-) with previously water-soaked dried filter paper for 2 min, then clear airflow for 30 s. They were transferred into a training tube, lined with sucrose or sucrose toxin mixture and exposed to a second odor (the CS+) for 2 min. To test immediate memory, flies were given clear airflow for 30 s then transferred to a T-maze and given 2 min to choose between the two odors. For 15, 30, 60 min and 6, 12, 18 and 24 h memory testing, flies were transferred back into starvation vials before being reloaded at the relevant time into the T-maze for testing. A Performance Index was calculated as the average score of two reciprocal experiments in which different populations of flies of the same genotype were trained to associate sucrose, or sucrose/toxin with either of the two odors. Odors used in conditioning were 8 μl 3-octanol (Sigma-Aldrich, 218405) diluted in 8 ml mineral oil (Sigma-Aldrich, 330760) and 13 μl 4-methyl-cyclohexanol (Sigma-Aldrich, 153095) diluted in 8 ml mineral oil. Airflow was maintained at 800 ml min^−1^. All experiments were performed at 23 °C and 50 % humidity unless otherwise stated.

For experiments using UAS-*dTrpA1*, flies were raised at 18 °C until starvation, which was performed at permissive 23 °C. Flies were then shifted (the timing of which differed depending on the respective experiment) to restrictive 32 °C for at least 15 min to activate neurons expressing *dTrpA1*. Similarly, for experiments involving UAS-*Shi^ts^*^1^, flies were kept at the permissive 23 °C until shifting to restrictive 32 °C for at least 20 min to block output of neurons expressing *Shi*^ts^^1^.

### b-glucosidase inhibitor feeding experiment

Flies were ad libitum fed Conduritol-beta-Epoxide (5mg in 500 ml of yellow food), an inhibitor of the b-glucosidase enzyme, for 4 days before starvation in preparation for malaise training.

### T-maze olfactory avoidance behavior

Olfactory avoidance behavior was assessed using a standard T-maze apparatus. Flies in the relevant conditioned state were introduced at the choice point and given a binary choice between two converging air streams. One arm carried a control stimulus consisting of air passed over 8 ml of mineral oil. The other arm carried an odorant stimulus: either octanol (OCT; 10 μl diluted in 8 ml mineral oil) or 3-methylcyclohexanol (MCH; 14 μl diluted in 10 ml mineral oil). An Avoidance Index (AI) was calculated as the difference between the number of flies choosing the control arm and the odorized arm: (Ncontrol−Nodor)/Ntotal, where Ncontrol represents the number of flies in the mineral oil control arm, Nodor represents the number of flies in the odorized arm, and Ntotal is the total number of flies tested in that trial. All assays were performed at 23 °C.

### Locomotor assays

Locomotion was measured using a modified protocol from Nojima et al ^134^. Individual flies were collected either 15 min or 24 h after malaise or sugar training. Assays were performed at 25 °C in custom-built round chambers (20 mm diameter × 2 mm height), each arena separated by a retractable divider. The chamber plates (4 × 5 arrangement, 20 chambers per plate) were fabricated from acrylic (Southern Acrylics, UK) and mounted on a custom 3D-printed support. Chambers were backlit with an FLFL-Si200-IR24 infrared backlight (Falcon Illumination). Videos were recorded using a Basler ace A2440-75um camera (cat. #35927) equipped with a 35 mm VIS–NIR fixed focal length lens (cat. #67716) and UV/VIS cut-off filter (cat. #89834, Edmund Optics). Recordings were captured as uncompressed AVI files at 2400 × 1600 resolution, 8-bit grayscale, 25 frames/s, for 15 min using LabVIEW software (National Instruments). Locomotor activity was quantified from video recordings using custom Python scripts.

### 2-Photon *in vivo* calcium imaging

All flies were raised at 25 °C and 3- to 5-day-old male and female flies were used in all experiments. Flies were briefly immobilized on ice and mounted in a custom-made chamber allowing free movement of the antennae and legs. The head capsule was opened under room temperature carbogenated (95% O_2_ and 5% CO_2_) buffer solution, and the fly, in the recording chamber, was placed under a two-photon microscope (Scientifica). For imaging fed flies, the following buffer was used: 103 mM NaCl, 3 mM KCl, 5mM N-Tris, 10 mM trehalose, 10 mM glucose, 7 mM sucrose, 26 mM NaHCO_3_, 1 mM NaH_2_PO_4_, 1.5 mM CaCl_2_ and 4 mM MgCl_2_, osmolarity 275 mOsm, pH 7.3). For acute 5-HT application, a perfusion pump system (14-284-201, Fisher Scientific) was used to continuously deliver saline at a rate of approximately 0.043 ml s^−1^. 5-HT (100 μM serotonin hydrochloride (H9523, Sigma Aldrich) was applied in the presence of 1 µM tetrodotoxin to block voltage-gated sodium channels and propagation of action potentials that could result in indirect excitation. To examine the effects of 5-HT on DANs, we acquired GCaMP7f fluorescence in targeted DANs such as PAM-β’2mp, PAM-γ3, PAM-γ5, PPL1-γ1pedc or PPL1-γ2α’. During pre-perfusion only saline was applied, and 5-HT was used during perfusion. Concentration of 5-HT used is comparable to recent physiological studies applying exogenous 5-HT to the *Drosophila* brain ^102–105^. To analyze two-photon fluorescence images were manually segmented using Fiji^106^, using custom-made Fiji script including an image stabilizer plugin^107^. Movement of the animals was small enough for images to not require registration. For subsequent quantitative analyses, GraphPad Prism (10) was used. The mean intensity of the GCaMP7f signal was measured using ImageJ (Fiji). For each brain, the average intensity of GCaMP7f signal of the 600 frames acquired during pre-perfusion was used as the baseline (F0). ΔF/F0 was calculated as [(average intensity of GCaMP7f signal of the 600 frames during perfusion) – (average intensity of GCaMP7f signal of the 600 frames during pre-perfusion)] / (average intensity of GCaMP7f signal of the 600 frames during pre-perfusion). Each *n* corresponds to a recording from a different individual fly.

For TotA perfusion, a synthetic predicted bioactive TotA peptide (RLYPSLTPEERESIDKF) was applied in the presence of 1 µM tetrodotoxin, Peptides were synthesized by GenScript.

For MBON imaging, flies trained in T-mazes using the malaise protocol were imaged at different time points afterwads using 2 photon microscopy. A constant air stream, carrying vapor from mineral oil solvent (air) was applied. GCaMP responses to the CS+, the CS- and a third odor were then measured in the relevant MBONs. Flies were sequentially exposed to the CS+, CS- and a third odor, isopentyl acetate (IAA; 1:10^3^ concentration) for 5 s. Each odor presentation was followed by 30 s of air. A plane through the dendritic field of each targeted MBON was selected for imaging. Fluorescence was excited using ∼140 fs pulses, 80 MHz repetition rate, centered on 910 nm generated by a Ti-Sapphire laser (Chameleon Ultra II, Coherent). Images of 256 × 256 pixels were acquired at 5.92 Hz, controlled by ScanImage 3.8 software ^135^. Odors were delivered using a custom-designed system ^136^. For analysis, two-photon fluorescence images were manually segmented using Fiji^106^. Movement of the animals was small enough such that images did not require registration. For subsequent quantitative analyses, custom Fiji and Python scripts were used. The baseline fluorescence, F_0_, was defined for each stimulus response as the mean fluorescence F from 2 s before and up to the point of odor presentation. F/F_0_ accordingly describes the fluorescence relative to this baseline. For the MBON imaging, the area under the curve (AUC) was measured as the integral of F/F_0_ during the 5 s odor stimulation.

### Transcriptional profiling

Serotonin receptor subtype expression in DANs innervating the MBs were measured using a combination of single-cell transcriptomes from several previous studies ^49, 112, 113^ and new data (Treiber et al., in preparation). Midbrain-wide transcriptomes from individual experiments were first aligned with each other and clustered using the Seurat v3 package^114^. Cell clusters expressing *Vmat*, encoding the monoamine vesicular transporter, and *pale* encoding tyrosine hydroxylase, were then extracted and reclustered. Clusters of these data were assigned to their corresponding anatomically defined DAN cell types using a combination of known marker genes ^115^ and the relative number of cells contributing to each cluster (Treiber et al., in preparation).

### Hemolymph collection

Hemolymph was collected using a centrifugation-based method. First, three small holes were pierced into the cap of a 0.5 mL Eppendorf tube. This pierced tube was then placed inside an open 1.5 mL Eppendorf tube to act as a collection chamber. Adult flies were anesthetized on ice, and their wings were removed. Each fly was immobilized by inserting a fine stylus through the thorax, and ∼20 prepared flies were placed into the pierced 0.5 mL tube. The nested tubes were centrifuged at 5000 rpm for 1 min at 4 °C. During centrifugation, hemolymph drained through the holes in the inner tube and collected at the bottom of the 1.5 mL tube. After centrifugation, the 0.5 mL tube containing the flies was discarded. Hemolymph in the 1.5 mL tube was then collected with a calibrated capillary tube, and the volume was recorded. On average, approximately 50 flies yielded ∼0.5 μL of hemolymph. The hemolymph could subsequently be transferred to other containers to store at −80 °C for further use.

### Proteomics: performed commercially MS sample preparation

Samples were lysed using 50 uL of lysis buffer (consisting of 6M Guanidinium Hydrochloride, 10 mM TCEP, 40 mM CAA, 50 mM HEPES pH8.5). The samples were boiled at 95C for 5 mins, sonicated on high for 5×60 sec on/30sec off using the bioruptor pico sonication water bath (Diagenode). Lysed samples were centrifuged at 10000*g for 10 mins, and supernatants were transferred to clean LoBind Eppendorf tubes. Protein concentration was determined by BCA rapid gold (Thermo) and 20μg of protein was taken forward for digestion. Samples were diluted 1:3 with digestion buffer (10% Acetonitrile in 50mM HEPES pH 8.5) and incubated with 1:100 enzyme to protein ratio of LysC (MS Grade, Wako) at 37C for 4 hours while shaking at 1000rpm. Samples were further diluted to a final 1:10 with more digestion buffer and digested with 1:100 trypsin (MS grade, Sigma) for 18 hours at 37C, shaking at 1000rpm.Samples were acidified by adding 2% trifluoroacetic acid (TFA) to a final concentration of 1%.Prior to TMT labeling, the peptides were desalted on a SOLAμ SPE plate (HRP, Thermo). Between each application, the solvent was spun through by centrifugation at 1500 RPM. For each sample, the filters were activated with 200ul of 100% Methanol, then 200ul of 80% Acetonitrile, 0.1% formic acid. The filters were subsequently equilibrated 2x with 200ul of 1% TFA, 3% Acetonitrile, after which the sample was loaded.

After washing the tips twice with 200ul of 0.1% formic acid, the peptides were eluted into clean 0.5ml Eppendorf tubes using 40% Acetonitrile, 0.1% formic acid. The eluted peptides were concentrated in an Eppendorf Speedvac and re-constituted in 20uL 50mM HEPES (pH8.5) for TMT labelling with 18plex tags (Thermo). A reference sample was prepared by mixing equal amounts of peptides from each sample and labeling that separately. Labeling was done according to manufacturer’s instructions, and subsequently, labeled peptides were mixed 1:1:1:1:1:1:1:1:1:1:1. TFA was added to acidify and bring acetonitrile concentration down to < 5%. Prior to mass spectrometry analysis, the peptides were desalted and fractionated using an offline ThermoFisher Ultimate3000 liquid chromatography system using high pH fractionation (5mM Ammonium Bicarbonate, pH 10) at 5ul/min flowrate. 10ug of peptides were separated over a 120min gradient (5% to 35% Acetonitrile), while collecting fractions every 130sec. The resulting 60 fractions were pooled into 30 final fractions, acidified to pH < 2 with 1% TFA and loaded onto EvoSep stagetips according to manufacturer’s protocol.

### MS data acquisition

For each fraction, peptides were analysed using the pre-set ’30 samples per day’ method on the EvoSep One instrument. Peptides were eluted over a 44-min gradient, and analysed with an Orbitrap EclipseTM TribridTM instrument (Thermo Fisher Scientific) with FAIMS ProTM Interface (ThermoFisher Scientific) switched between CVs of −50 V and −70 V with cycle times of 1.5 s. Full MS spectra were collected at a resolution of 120,000, with normalized AGC target set to 100% or maximum injection time of 50 ms and a scan range of 375–1500 m/z. MS1 precursors with an intensity of >5×103 and charge state of 2-7 were selected for MS2 analysis. Dynamic exclusion was set to 120 s, the exclusion list was shared between CV values and Advanced Peak Determination was set to ‘off’. The precursor fit threshold was set to 70% with a fit window of 0.7 m/z for MS2. Precursors selected for MS2 were isolated in the quadrupole with a 0.7 m/z window. Ions were collected for a maximum injection time of 35 ms and normalized AGC target set to ‘custom’. Fragmentation was performed with a HCD normalised collision energy of 30% and MS2 spectra were acquired in the IT at scan rate rapid. Precursors were subsequently filtered with an isobaric tag loss exclusion of TMT and precursor mass exclusion set to 25 ppm low and 25 ppm high.

Precursors were isolated for an MS3 scan using the quadrupole with a 2 m/z window, and ions were collected for a maximum injection time of 86 ms and normalized AGC target of 300%. Turbo TMT was deactivated and number of dependent scans set to 10. Isolated precursors were fragmented again with 50% normalised HCD collision energy, and MS3 spectra were acquired in the orbitrap at 50000 resolutions with a scan range of 100-500 m/z. MS performance was verified for consistency by running complex cell lysate quality control standard.

### Proteomics data analysis

The raw files were analysed using Proteome Discoverer 2.4 (Thermo Fisher Scientific). TMT reporter ion quantitation was enabled in the processing and consensus steps, and spectra were matched against the Drosophila melanogaster database (taxonomy 7227) obtained from UniProt. Dynamic modifications were set as Oxidation (M), and Acetyl on protein N- termini. Cysteine carbamidomethyl (C) and TMT 16-plex (peptide N-termini and K) were set as static modifications. All results were filtered to a 1% FDR, and protein quantitation done using the built-in Minora Feature Detector. The data was filtered so only proteins with 2 unique peptides and at least 60% valid values across all of the samples, or at least 75% within an experimental group were used in the downstream data analysis. The data was normalized using the “scale” method from limma ^137^. Statistical significance was assessed by either limma or wilcoxon signed-rank test depending on whether the proteins demonstrated a normal distribution or not. Filtration and statistical analysis was performed in the R programming language.

### Statistics and Reproducibility

Flies in an experimental group were selected randomly. Experimental and control groups were tested in random orders. No blinding was used in data collection and analysis. All repeats in this study were biological replicates (that is, different groups of flies in behavioral experiments and different fly brains in imaging experiments) and they were performed on at least two different days with similar results. Statistical analysis was performed in Prism 8 & 10. All behavioral data were tested for normality using the Shapiro–Wilk normality test, and no data was excluded from the analyses. To compare more than two groups, normally distributed datasets were analyzed using one-way ANOVA and Tukey’s multiple comparisons test. Data that were not normally distributed were analyzed using Kruskal– Wallis and Dunn’s multiple comparisons test. When comparing two groups, we used a two- tailed unpaired Student’s *t*-test with Welch’s correction for unpaired data or a two-tailed paired Student’s *t*-test for paired data for normally distributed data. Paired data that were not normally distributed were analyzed using a two-tailed Wilcoxon matched-pairs signed rank test. No statistical methods were used to predetermine sample sizes, but our sample sizes are equivalent to those in previous publications ^46,47^.

## Notes

### Competing Interest Statement

The authors have declared no competing interest.

