## Supplementary figures and images for "Visceral signaling of post-ingestive malaise directs memory updating in Drosophila"

### Figure S1

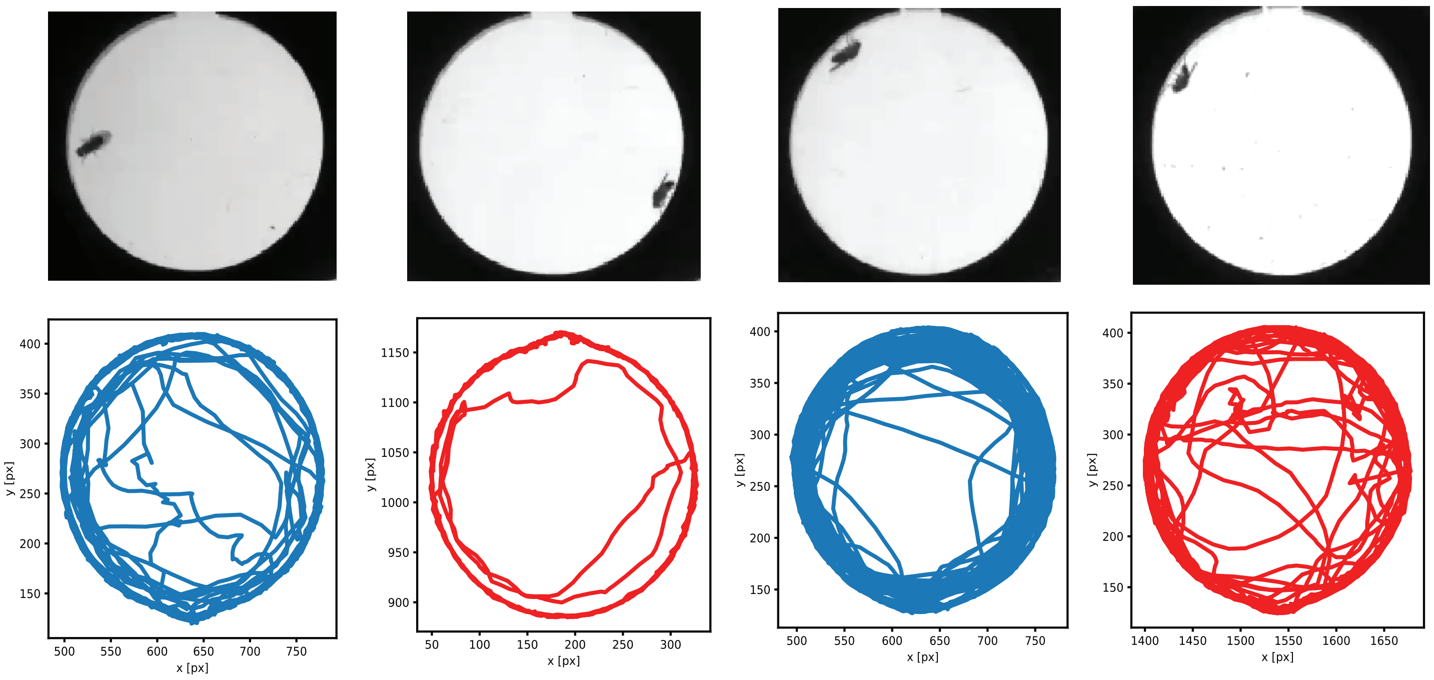

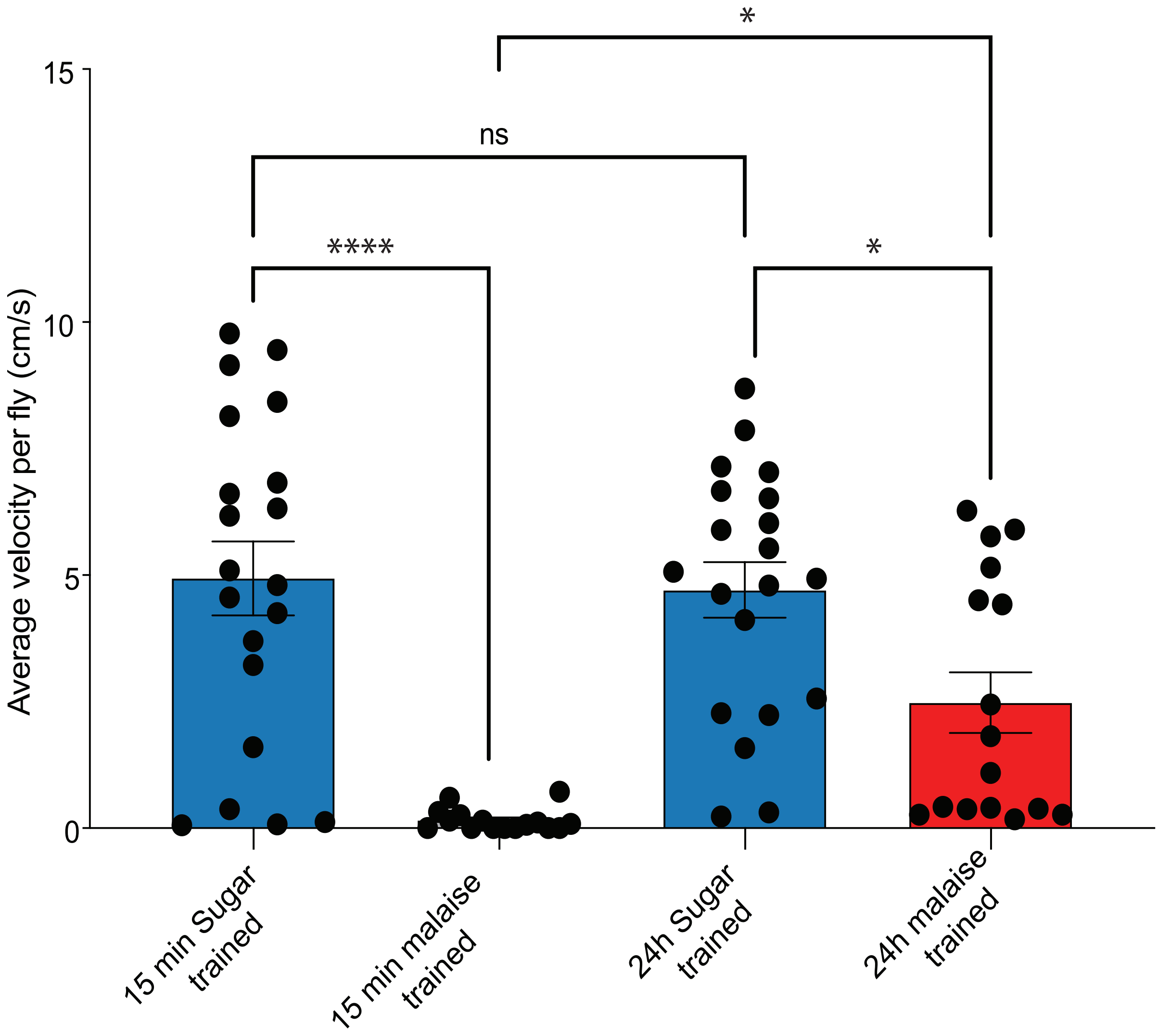

### Figure S2

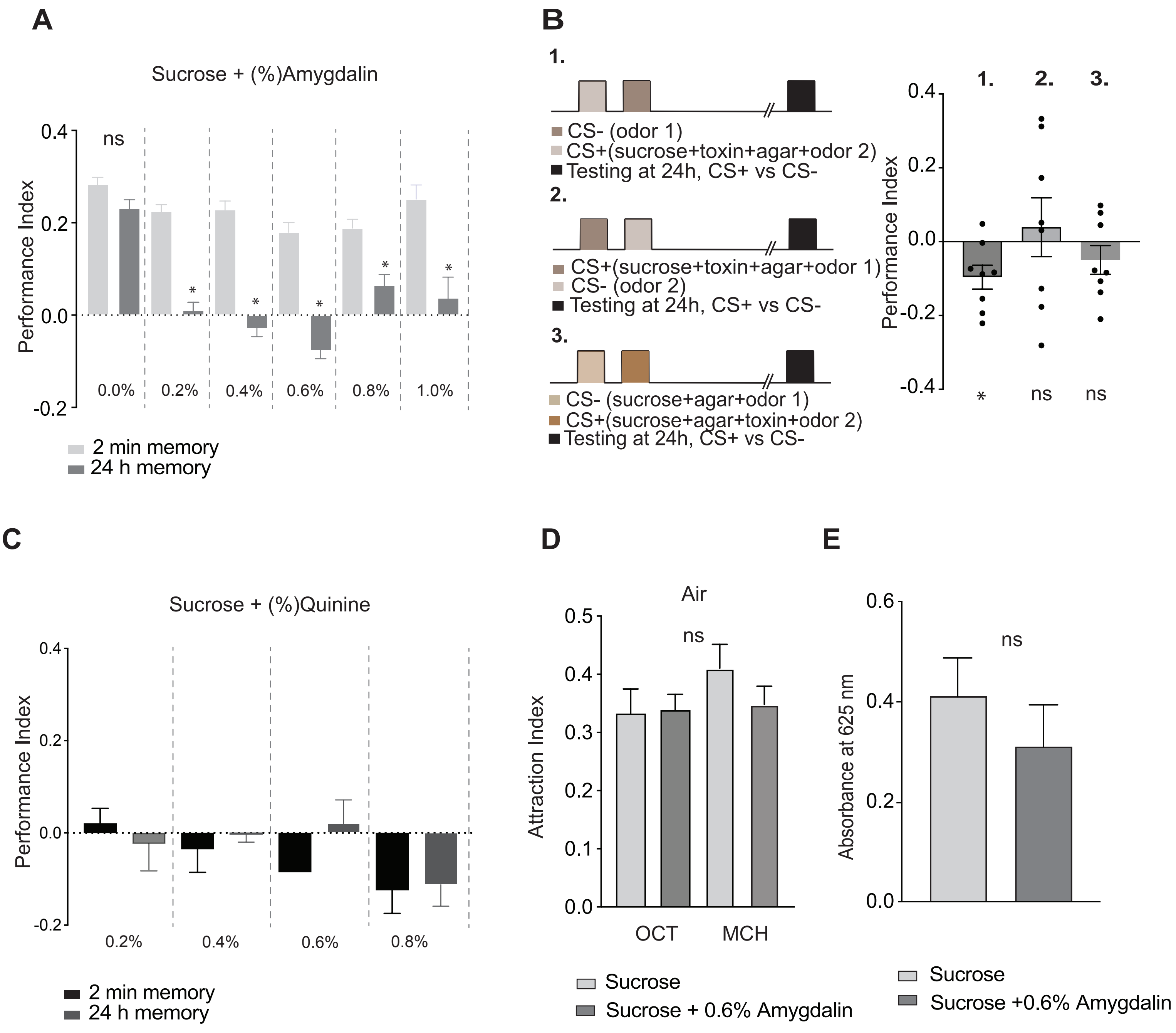

### Figure S3

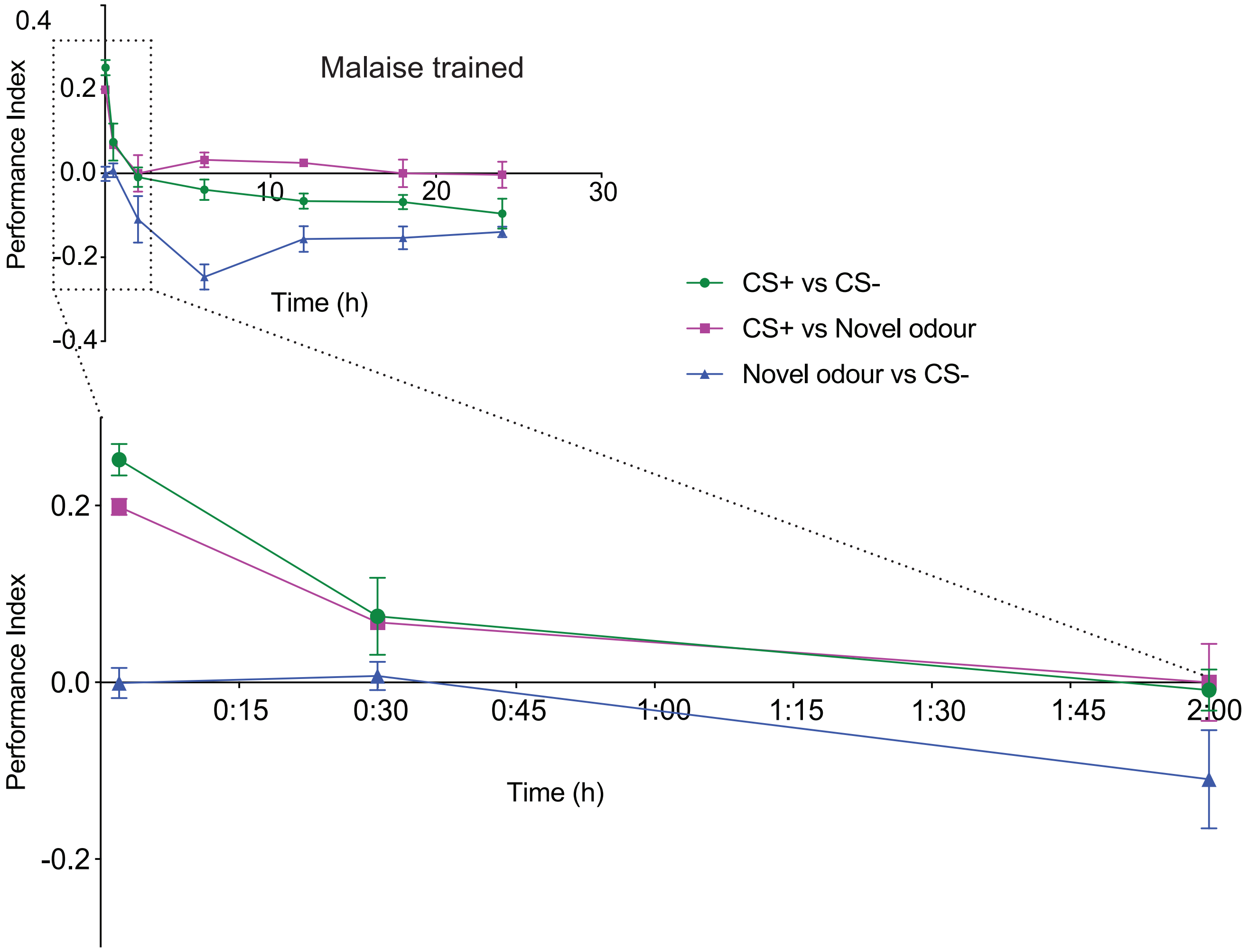

### Figure S4

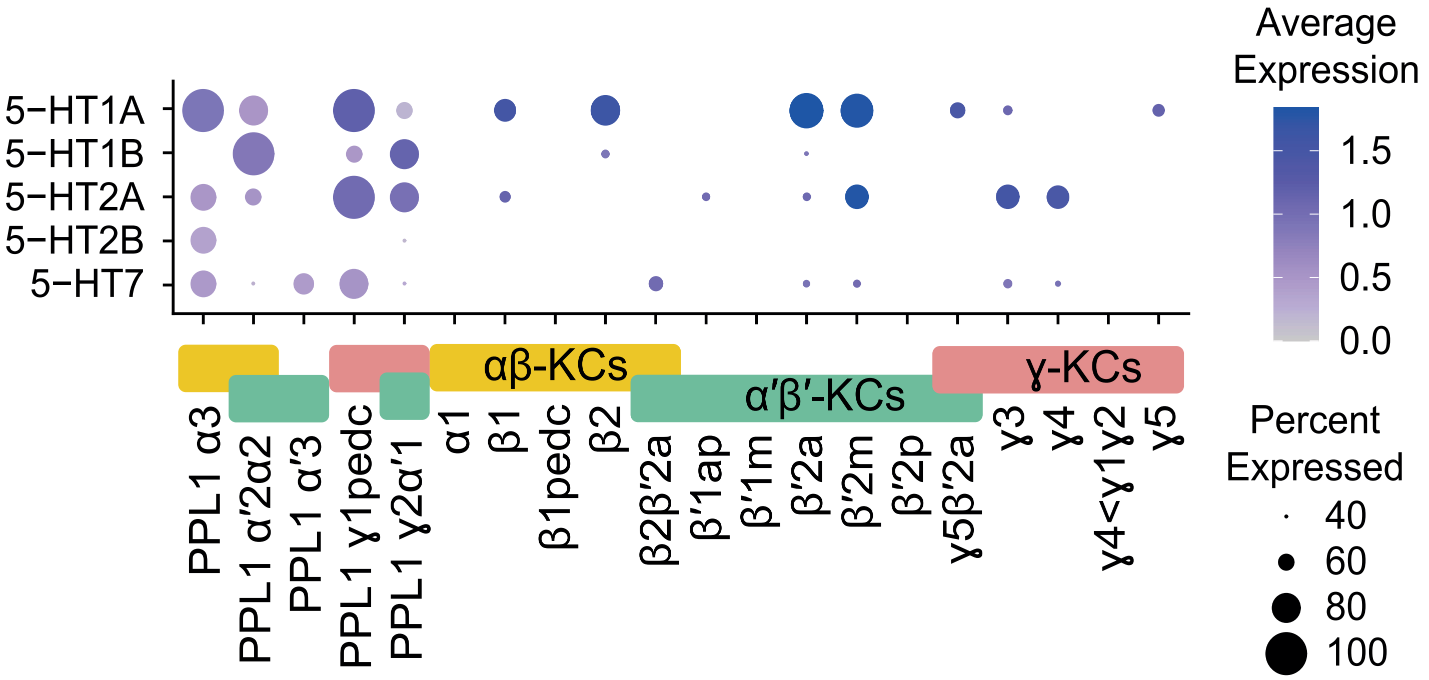

### Figure S5

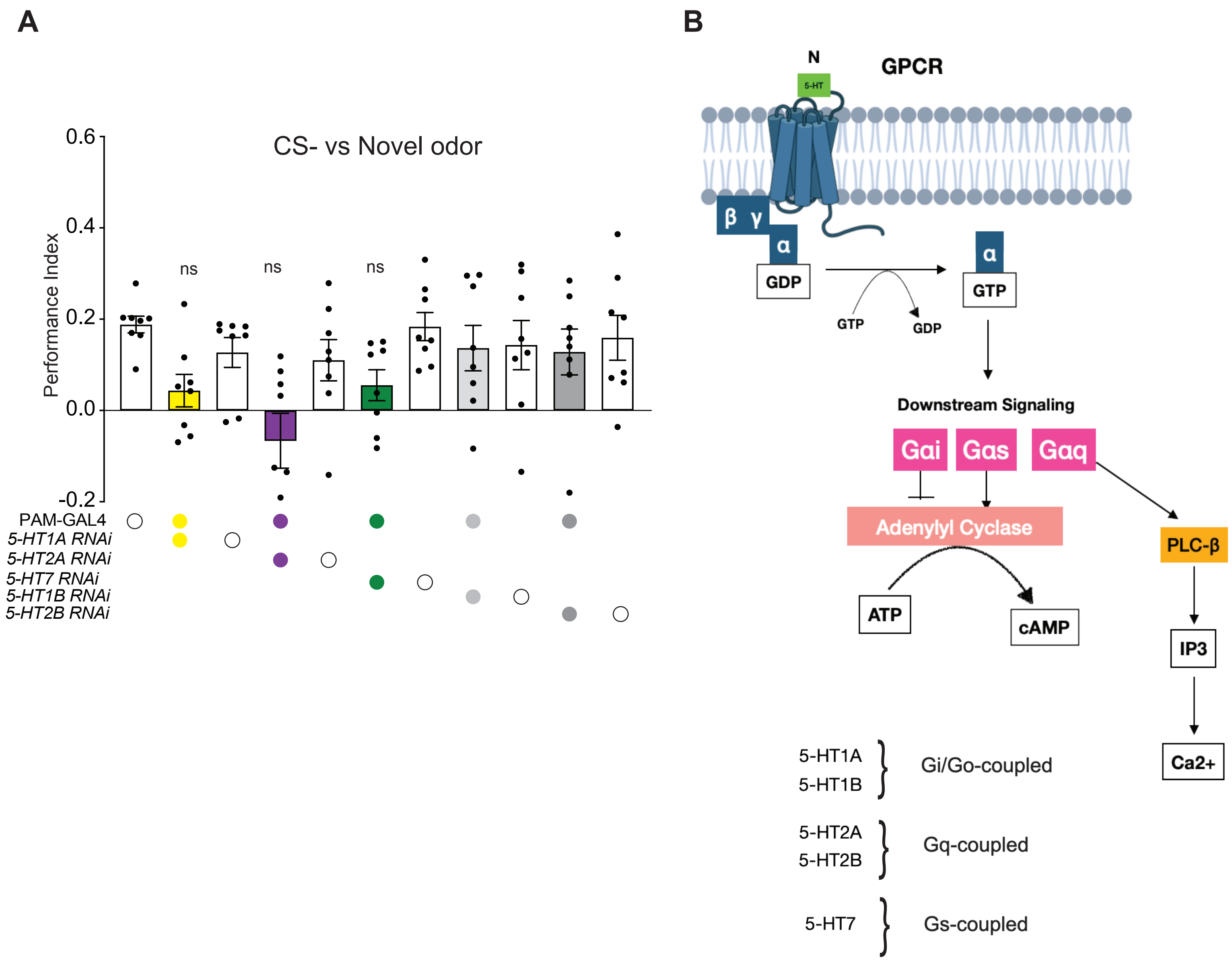

### Figure S6

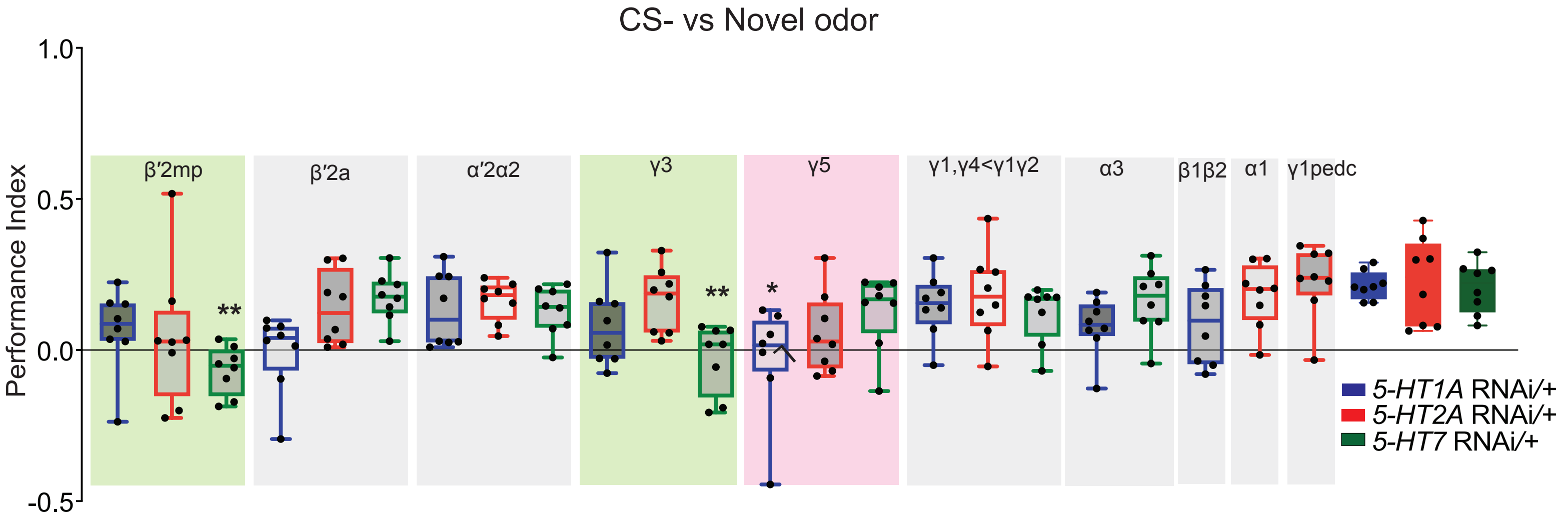

### Figure S7

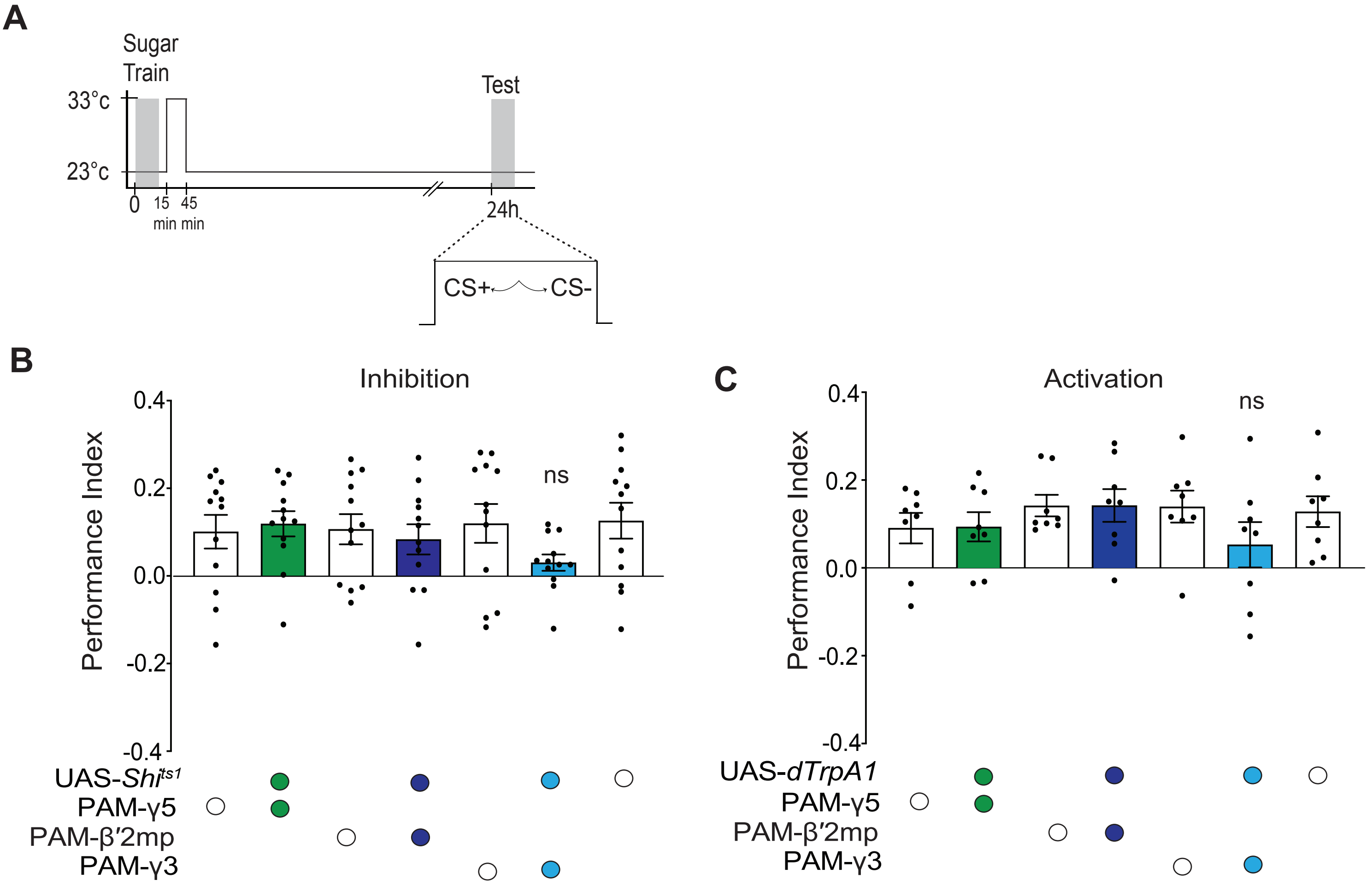

### Figure S8

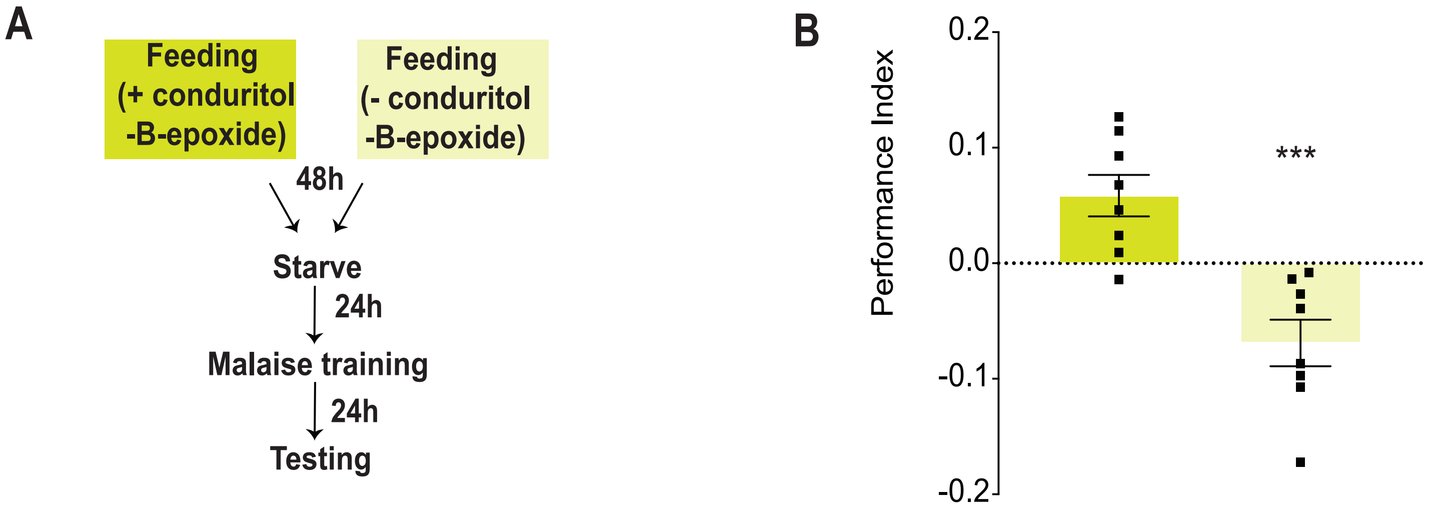

### Figure S9

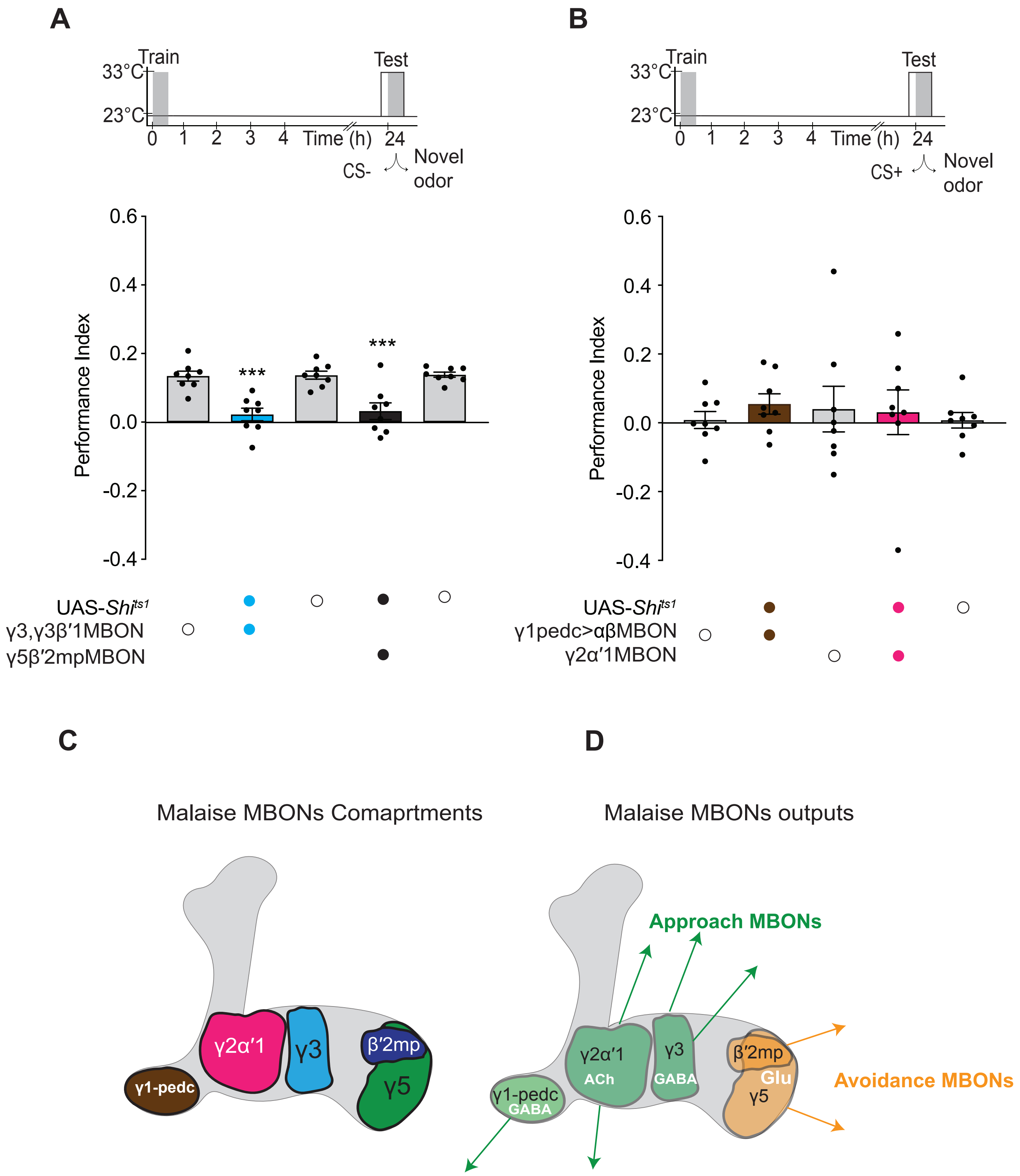

### Figure S10

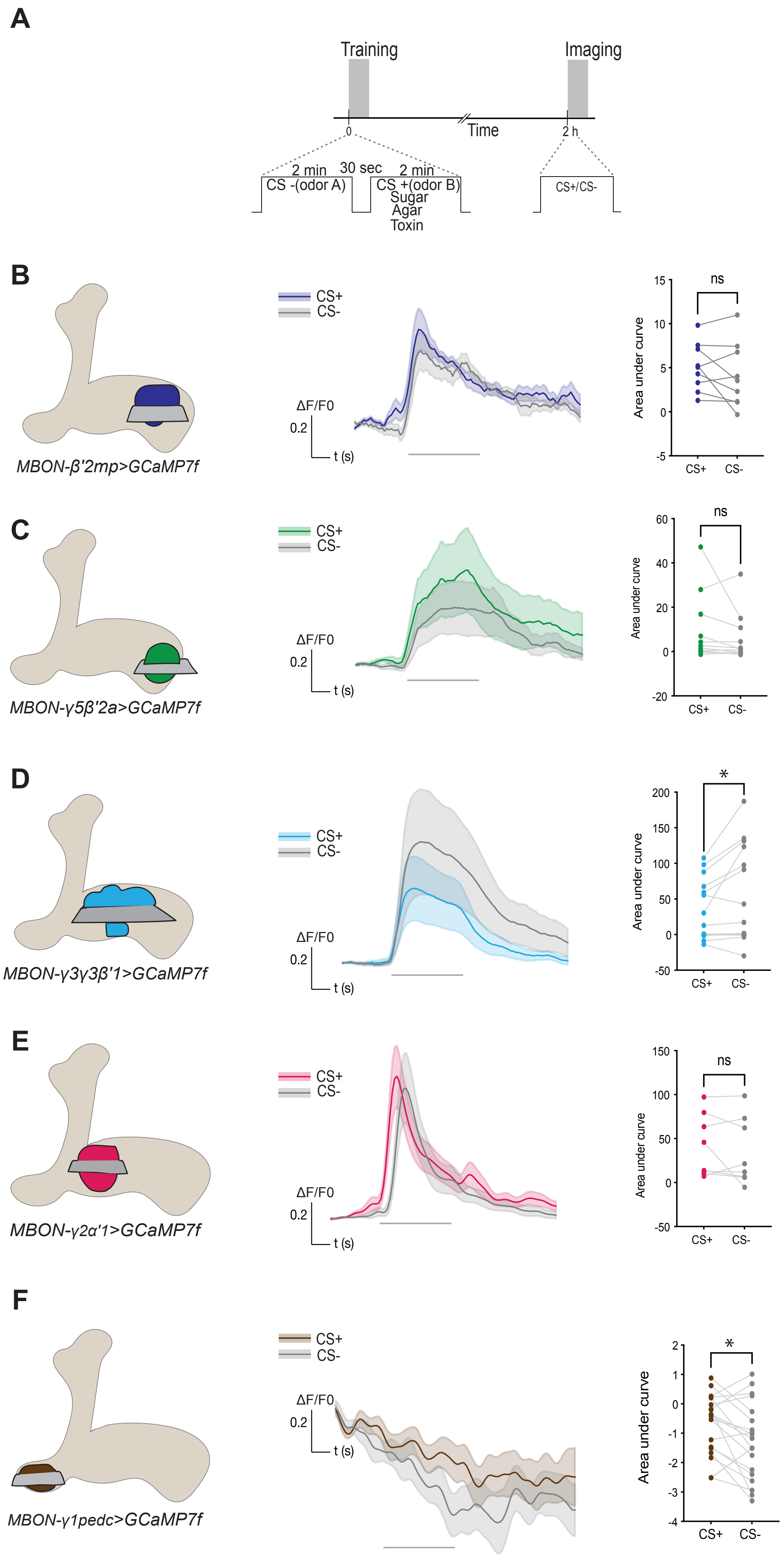

### Figure S11

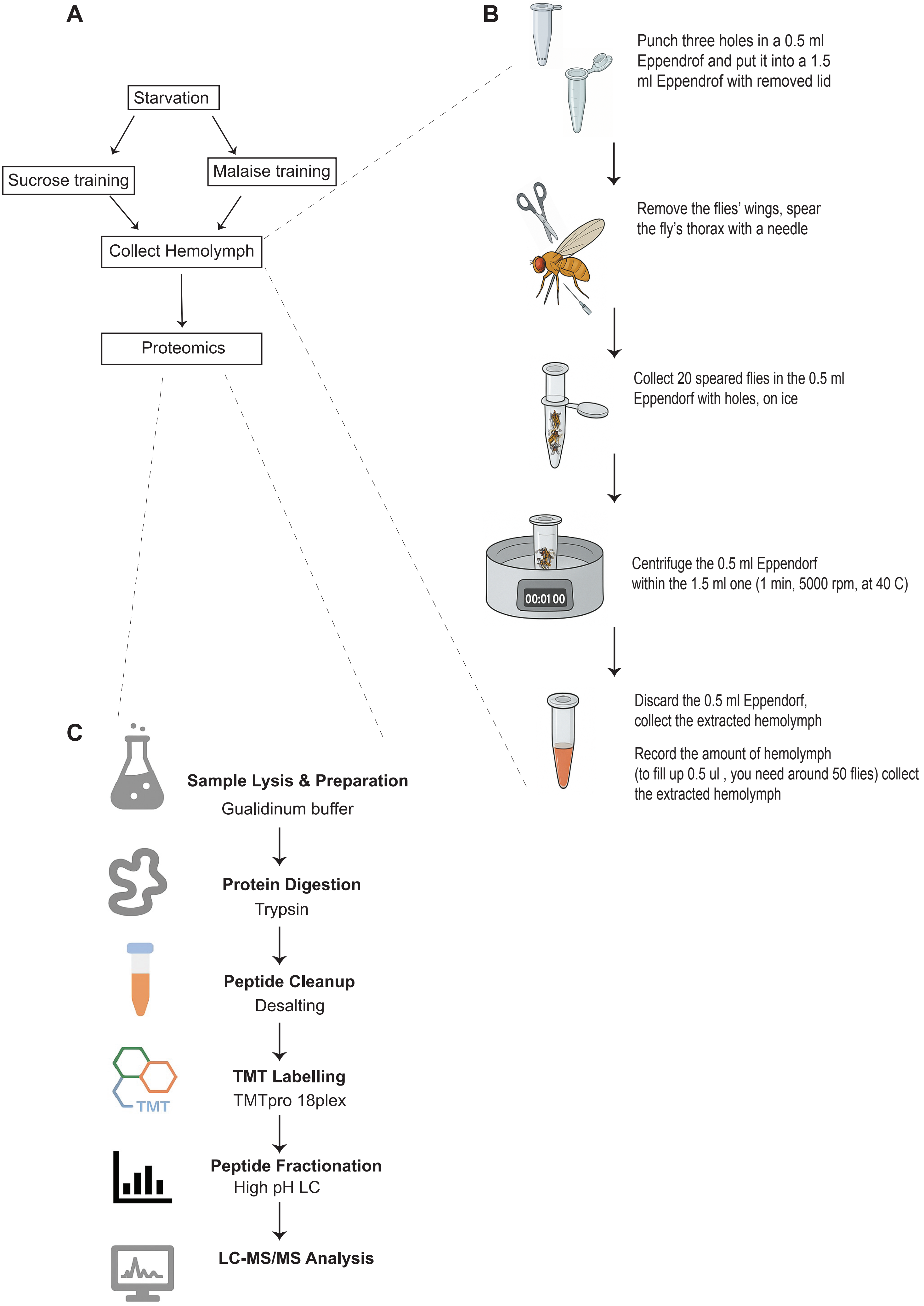

### Figure S12

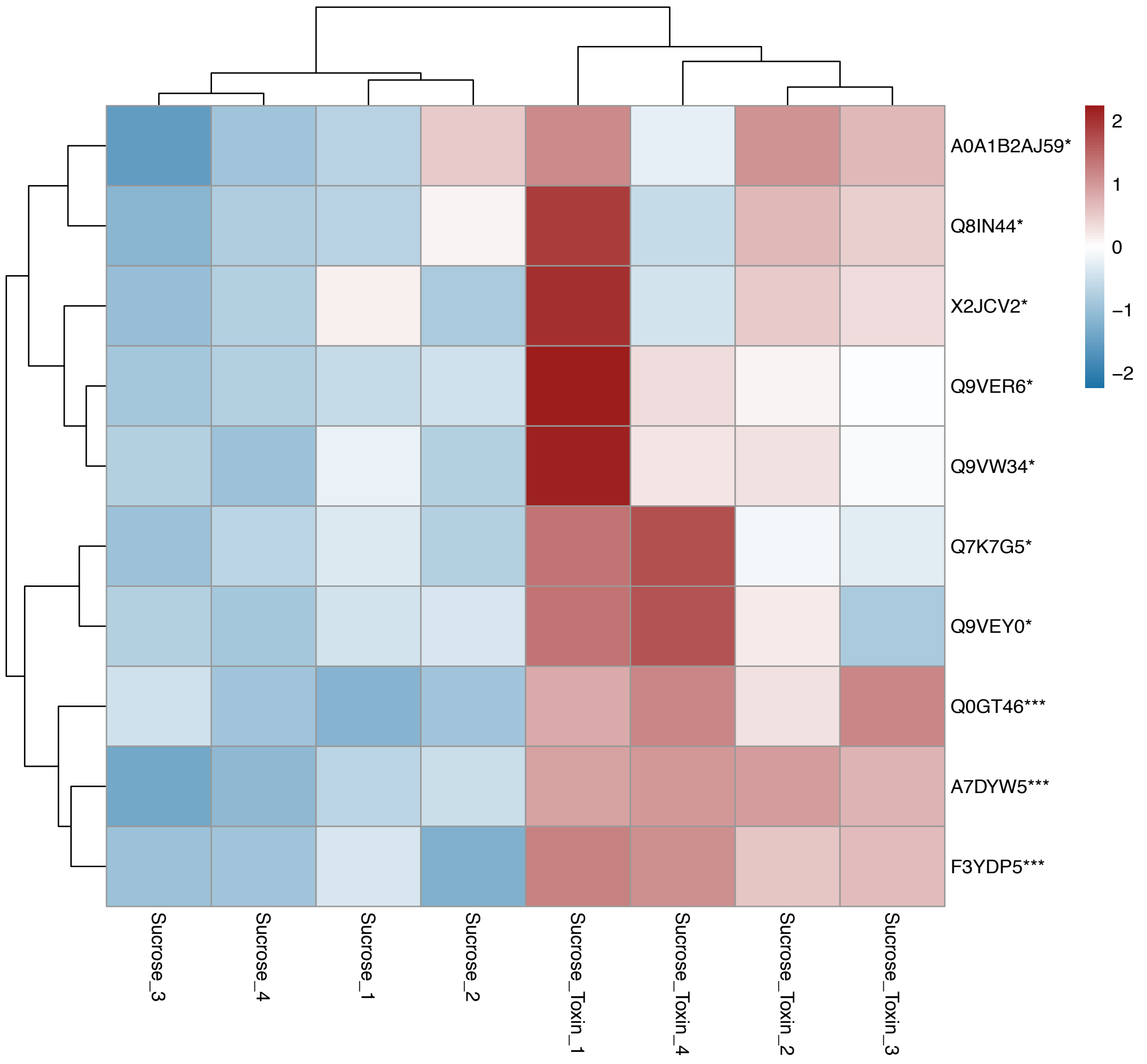

### Figure S13

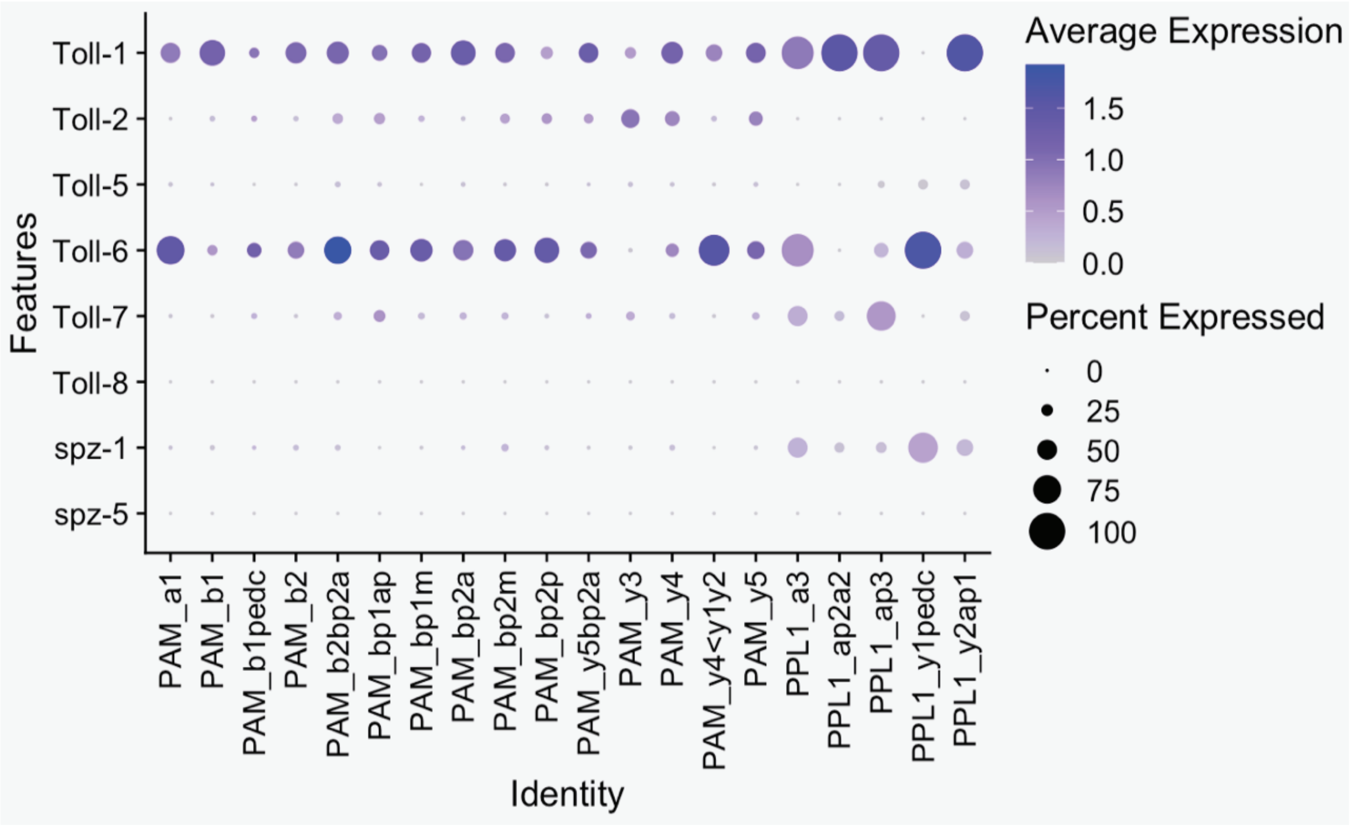
